# Dorsal striatal circuit mechanisms contributing to astrocyte modulation of alcohol-related behaviors

**DOI:** 10.64898/2026.06.01.729412

**Authors:** Cherish Ardinger, Anagha Kalelkar, Maxwell Madden, Ananya Gunda, Arnav Patel, George Xanthos, Mariam Mahboob, Ali Khawaja, Nancy Collie-Beard, Miriam Bocarsly, Rafiq Huda

**Author notes:** Corresponding author: Rafiq Huda.

## Abstract

**Background:** The dorsolateral striatum (DLS) is a key site for coordinating the alcohol-induced stimulant response, a behavioral marker predictive of future alcohol use disorder. Although ethanol (EtOH) affects all brain cells, little is known about the contribution of non-neuronal DLS cell types to EtOH-induced stimulation.

**Methods:** We used ex vivo two photon calcium imaging, in vivo fiber photometry of astrocyte and neuronal GCaMP, and astrocyte-specific manipulations in mice to determine DLS astrocyte contributions to EtOH-induced stimulation and voluntary EtOH drinking behavior. Using fiber photometry of GRAB-ACh sensors and cell-type specific chemogenetics, we also assessed the role of cholinergic signaling in observed astrocyte EtOH effects.

**Results:** As expected, intraperitoneal EtOH injections (0.5-2g/kg) evoked a stimulant response, evidenced by increased locomotion compared to saline. In parallel, EtOH dose-dependently decreased astrocyte calcium activity but had minimal effects on direct and indirect pathway neuronal activity. Mimicking this reduction with astrocyte-specific expression of CalEX, a calcium extruding pump, facilitated EtOH stimulation compared to mice expressing a control fluorophore. Hence, EtOH-induced suppression of DLS astrocyte activity contributes to stimulation. Astrocyte calcium signaling is a well-known target of neuromodulation. Fiber photometry recordings of extracellular acetylcholine (ACh) levels via GRAB-ACh imaging showed inhibition of ACh release by acute EtOH. We virally expressed the excitatory chemogenetic actuator hM3Dq in striatal cholinergic interneurons to assess whether artificially increasing ACh release blocks EtOH-induced inhibition of astrocytic calcium activity. Despite facilitating ACh release, this manipulation did not impact astrocyte calcium activity under control (saline) or EtOH conditions. Together, this work identifies DLS astrocytes as key contributors to EtOH-induced stimulation and highlights the importance of considering astrocyte-neuron interactions in evaluating alcohol effects.

## Introduction

AUD is a multi-faceted disorder with few FDA-approved treatment options available. Current FDA-approved pharmacotherapies for AUD target neuronal mechanisms to reduce alcohol intake [1]. However, alcohol impacts neurons and non-neuronal glia cells, including astrocytes [2]. Astrocytes have non-overlapping spatial territories and express a variety of neurotransmitter receptors which make them both anatomically and functionally capable of responding to synaptic transmission and neuromodulation with intracellular calcium elevations [3]. In turn, astrocyte calcium activity regulates diverse functions like neurotransmitter uptake and ionic homeostasis, neurovascular coupling, and direct neuronal modulation through release of bioactive substances [4, 5]. The fine processes of a single astrocyte can contact thousands of neuronal synapses, making astrocytes uniquely positioned to modulate synaptic transmission and integration [6, 7]. Growing evidence suggests that astrocyte calcium signaling plays a role in regulating behavior by modulating communication between ensembles of neurons [8–10]. Importantly, astrocytes also modulate alcohol-related behaviors. For example, cortical astrocytes regulate alcohol intake in mice [11] and activation of astrocytes in the basolateral amygdala decreases binge-like drinking [12]. Further, astrocytes in the nucleus accumbens have been implicated in behavioral flexibility after chronic alcohol exposure [13]. Despite these advancements, how alcohol affects in vivo astrocyte calcium activity and the underlying mechanisms remain largely unclear.

Alcohol produces divergent and heterogenous behavioral responses across individuals [14, 15]. Humans with heightened stimulant and decreased sedative responses to alcohol are at higher risk for developing alcohol use disorder (AUD) [16–21], and prevailing theory suggests a relationship between the initial stimulant response to alcohol and future alcohol intake [22, 23]. This subjective response to alcohol has recently been shown to also predict alcohol self-administration in heavy drinkers [24]. In rodents, alcohol-induced stimulation and sedation are commonly measured through the assessment of locomotor activity following injected alcohol [25–28]. In mice, the stimulant response to alcohol correlates with AUD-relevant behaviors like alcohol consumption and aversion-resistant drinking [25]. Relatedly, a lower sedative response correlates with alcohol consumption [29]. Like in humans, these preclinical findings show significant interindividual variability in alcohol behavioral sensitivity that correlates with AUD-relevant drinking behaviors. Hence, alcohol stimulation represents an avenue for reverse translation into pre-clinical models amenable to mechanistic dissection of neurobiological processes that jointly contribute to alcohol stimulation and associated AUD vulnerability.

Human neuroimaging studies and preclinical work both implicate the dorsolateral striatum (DLS) as an important regulator of alcohol’s stimulant effects [25, 30]. The DLS is a major basal ganglia input nucleus that integrates cortical and thalamic glutamatergic innervation under strong neuromodulatory control from midbrain dopaminergic neurons and local cholinergic interneurons [31–33]. Subgroups of striatal projection neurons (SPNs) differentially express dopamine D1 and D2 receptors, defining the direct and indirect pathways of the striatum, respectively. Although dopamine signaling in DLS SPNs modulates alcohol stimulation [25], the role of cholinergic signaling in this process is not known. Similarly, the contribution of astrocyte calcium activity to alcohol stimulation and sedation is not clear. Alcohol inhibits cerebellar and cortical astrocyte calcium activity via inhibition of norepinephrine (NE) release [34]. A similar mechanism in the DLS is unlikely given sparse NE innervation of the striatum. Astrocytes express multiple subtypes of cholinergic receptors, including GPCR coupled muscarinic and ionotropic nicotinic receptors, and respond to cholinergic neuromodulation with intracellular calcium increases [35–38]. In the current study, we use a range of ex vivo and in vivo approaches to determine how ethanol (EtOH) impacts astrocyte calcium signaling in the DLS and the relevance of this modulation for EtOH stimulation and consumption behaviors. We find that EtOH induced an expected locomotor stimulant response, along with an inhibitory effect on astrocyte calicum signaling and extracellular ACh levels in-vivo. Molecular genetic inhibition of astrocyte calcium signaling potentiated the EtOH-induced stimulant response, suggesting a role for DLS astrocytes in modulating this behavior. We also evaluated whether and how cholinergic activity modulates astrocyte alcohol effects. Together, our experiments highlight DLS astrocyte and cholinergic signaling as key coordinators of alcohol stimulation.

## Materials and methods

### Mice

All experimental procedures performed on mice were approved by the Rutgers University Institutional Animal Care and Use Committee (Protocol #202000004) and conformed with NIH guidelines. Progenitor mice were purchased from vendors (see Supplementary Table 1 for details), bred in-house, and used for experiments. A few experiments were performed with C57BL/6J mice purchased from Jax, which are noted in Supplementary Table 2. When appropriate, offspring were genotyped using Transnetyx services. All mice were at least 8 weeks of age prior to performing stereotaxic surgeries. Following surgery, mice were housed in a reversed 12-h light/dark cycle in standard-sized cages and provided ad libitum food (LabDiet 5R53) and water.

### Stereotaxic surgery

Viral infusions, head bar implants, and fiber optic implant surgeries were conducted as described by our lab previously [39, 40]. Specific viral infusion volumes and placements for each experiment can be found in Supplementary Table 2. For fiber optic surgeries targeting the dorsolateral striatum (DLS) and dorsomedial striatum (DMS), ceramic black fibers from RWD (907-03019-00), 400 µm core, 0.5 NA, 4 mm length were used. For fiber optic surgeries targeting the bed nucleus of the stria terminalis (BNST), the same fibers were used at a 5 mm length.

### Slice imaging using two photon microscopy

#### Brain slice preparation

C57BL/6 mice (n=6; 3M, 3F) received bilateral stereotaxic DLS (A/P: +0.3, ML: ± 2.5, DV: -3.3, [41]) injections of gfaABC1D-cyto-GCaMP6F. Following recovery and viral expression, mice were deeply anesthetized with isoflurane prior to decapitation. The brain was immediately removed and placed within an ice chilled cutting solution composed of 194 mM Sucrose, 30 mM NaCl, 4.5 mM KCl, 1 mM MgCl2, 26 mM NaHCO3, 1.2 mM NaH2PO4, 10 mM Glucose. 250 μm thick coronal slices containing the dorsal striatum were cut with a Leica VT1200 vibrating microtome. The vibratome chamber was ice-chilled prior to cutting and was filled with oxygenated cutting solution. Slices were incubated at 32 °C in oxygenated artificial cerebrospinal fluid (aCSF). The aCSF contained 124 mM NaCl, 4.5 mM KCl, 1 mM MgCl2, 26 mM NaHCO3, 1.2 mM NaH2PO4, 10 mM Glucose, and 2 mM CaCl2. The cutting solution and aCSF were prepared with deionized water (18.2 MΩ-cm), filtered (0.22 µm), and bubbled with 95% O2 and 5% CO2 for at least 15 min prior to use and throughout slice preparation and recording. After incubation, slices were maintained at room temperature until 2 photon calcium imaging, which was performed in the same aCSF formulation used for incubation.

#### Brain slice two photon calcium imaging & analysis

GCaMP6f fluorescence from astrocyte somas was imaged through a 16x/0.8 NA objective (Nikon) using resonance-galvo scanning with a Ultima Investigator two-photon microscopy system (Bruker). Image frames were 512 × 512 pixel resolution acquired at 64 Hz with 64-frame averaging resulting in an effective frame rate of 1 Hz. Emitted light was filtered using a dichroic mirror and collected with GaAsP photomultiplier tubes (Hamamatsu). GCaMP6f-expressing astrocytes were imaged with 2x optical zoom. Astrocyte soma fluorescence over time was extracted using ImageJ, and a ΔF/Fo (DFF) was calculated as

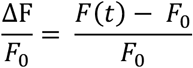

Where F0 represents the 8^th^ percentile of the fluorescence values recorded in the surrounding two minutes. The “scipy.signal.find_peaks” function was utilized to identify calcium events in the DFF trace with minimum peak prominence of 0.4 DFF. From the identified peaks, both inter event interval (IEI) and average peak amplitude were determined. All recordings were visually examined for drift in the imaging plane. Where such drift occurred transiently (<30 seconds) the affected period was excluded from analysis, in other cases the entire recording was excluded from analysis.

### Fiber photometry recordings of bulk astrocyte and neuronal calcium activity

We used fiber photometry to assess how acute EtOH impacts astrocytic calcium signaling during ethanol (EtOH)-induced stimulant response open field testing. Striatal astrocytes of C57BL/6J mice were unilaterally transfected with the genetically encoded Ca^2+^ indicator AAV-gfaABC1D-cyto-GCaMP6F (n = 7; 4F, 3M, DLS) or pZac2.1-gfaABC1D-lck-GCaMP6f (n = 11; 5F, 6M, [AP: +1.00, ML: ±1.5, DV: -2.6; DMS]). See Supplementary Table 1 for detailed information for each construct. Calcium activity from cell-type specific striatal neurons was recorded using pGP-AAV-Syn-FLEX-jGCaMP8m-WPRE stereotaxically injected into the DLS of transgenic mice (Drd1-Cre: n = 5; 3F, 2M; A2a-Cre: n = 6; 4F, 2M). For all fiber photometry experiments, at least 3 weeks was given following surgery to allow for viral expression. Fiber recordings were conducted using the Tucker-Davis Technologies system as described by our lab previously [40].

#### Fiber photometry analysis

Fiber photometry data were analyzed using custom MATLAB scripts, as previously described by our lab [40]. Briefly, raw fiber photometry data was imported using the opensource file “TDTbin2mat.m,” which is available for download on the TDT website. Data from emission at 465 nm (signal) and 405 nm (isobestic control) were both down sampled to 20 Hz. Then a linear fit of the isobestic 405 nm channel to the 465 signal was calculated using the ‘polyfit’ MATLAB function. Lastly, a commonly used ΔF/F calculation was performed to control for photobleaching and movement artifacts, where ΔF/F = (signal – fitted control) / fitted control. Event rate and event prominence were calculated from these ΔF/F traces using the ‘findpeaks’ MATLAB function.

### EtOH-induced open field stimulant response testing

Open field testing was conducted using a 40 × 40 × 40 cm arena (60101, Stoelting). For all open-field tests, mice received 3 days of habituation where they received an intraperitoneal injection of sterile 0.9% saline and were allowed to acclimate to the open-field arena for 30 minutes. Mice were then randomly assigned to receive one dose of EtOH prior to testing (saline, 0.5, 1, 1.5, or 2 g/kg) on a given day. EtOH was injected at a volume of 10 mL/kg (i.e., for 0.5 g/kg EtOH the concentration was 6.25% v/v; for 1 g/kg it was 12.5% to ensure injection volumes were consistent with body weight across testing days regardless of dose). Saline injections were volume-matched with EtOH injections. Note that doses were given in a randomly assigned order and mice only received one EtOH injection per day to allow them to metabolize the dose completely before the next testing session, which typically occurred as subsequent testing days. EtOH solutions were prepared from 200 proof EtOH (Koptec) diluted in sterile 0.9% saline. An overhead camera (Microsoft) captured open field video to assess movement and fiber recordings were conducted as described above if applicable. For astrocytic and neuronal calcium recordings (Figures 1 and 3) and CalEX open field experiments (Figure 2), mice were injected with EtOH and immediately placed into the center of the open field for 30 minutes of testing. For neuromodulator recordings, a 10-minute baseline fiber recording conducted in the open field preceded the EtOH injection. For DREADD stimulation of cholinergic neurons, a drug pre-treatment of Clozapine-N-oxide (CNO; HelloBio) or Deschloroclozapine (DCZ; Sigma Aldrich) and 10-minute baseline fiber recording preceded EtOH injection (described in “Chemogenetic manipulation of cholinergic interneurons (CINs)” in more detail below).

**Figure 1.**
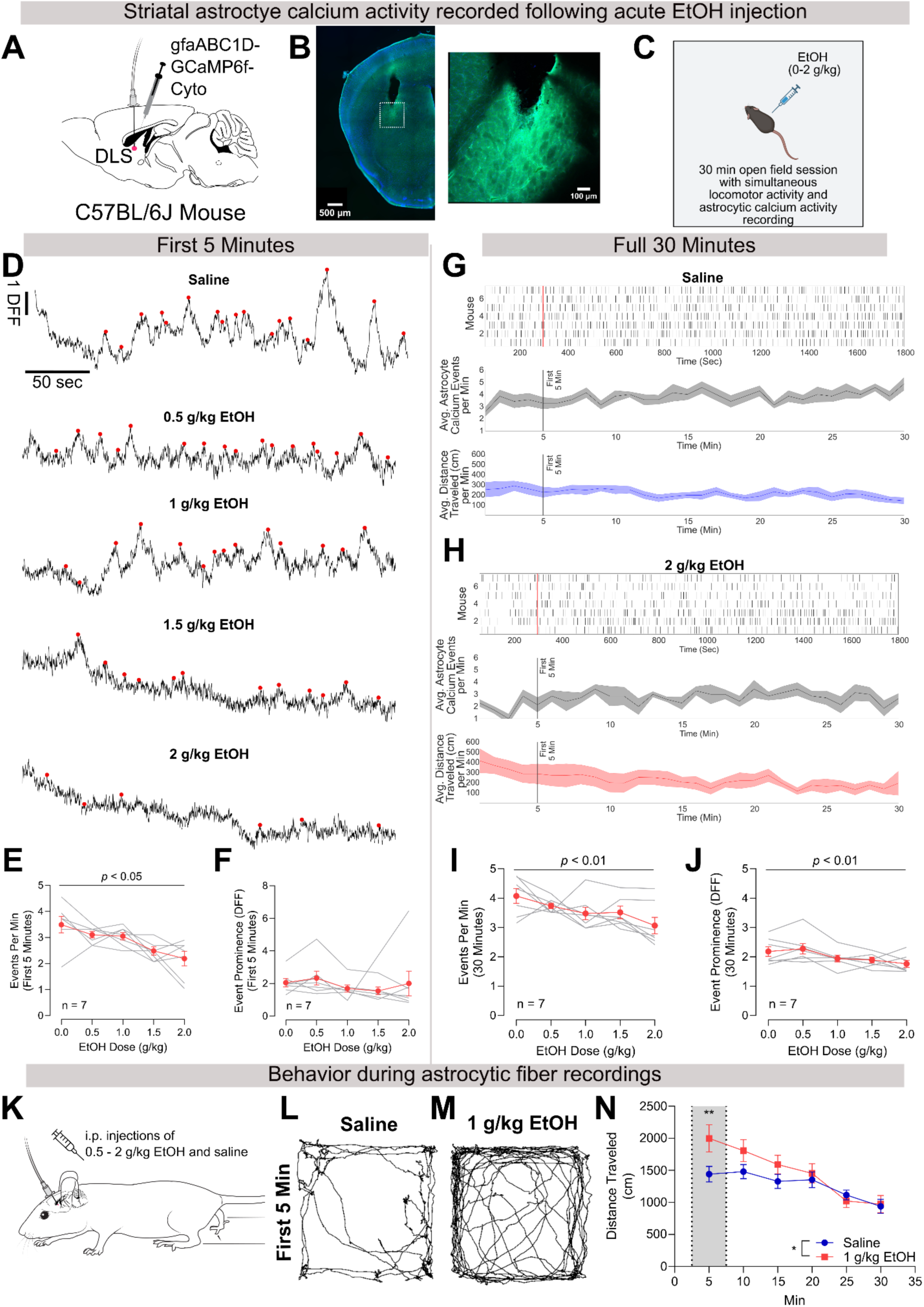
Alcohol acutely inhibits DLS astrocytic calcium signaling. **A)** Mice were given a stereotaxic viral injection of GCaMP6f with an astrocyte-specific promoter along with an optical fiber to enable in-vivo fiber photometry recordings. **B)** A representative image showing the DLS fiber placement and GCaMP6f viral expression (left) and zoomed-in image of the GCaMP6f expression (right). The dotted box on the left image represents the zoomed-in portion shown on the right image. **C)** Mice received saline or 0.5 – 2 g/kg EtOH injected intraperitoneally and then were placed in an open field box for 30 minutes while overhead video was recorded to assess locomotor activity simultaneously with astrocyte calcium activity gathered via fiber photometry. This panel was created using Biorender. **D)** Representative traces of astrocyte calcium activity from the first 5 minutes of an open-field recording across all doses tested. Red dots represent detected events. Note that all traces are from the same mouse and the scale bars from saline apply to all doses. **E)** In the first 5 minutes, alcohol inhibited astrocyte calcium events; *F*(2.380, 17.26) = 5.518, *p* = 0.011 (Greenhouse-Geisser corrected one-way repeated measures mixed-effects analysis). Tukey’s multiple comparison post-hoc tests indicated no significant differences between any two given doses; but a trend was identified in the post-hoc comparison of saline to 2 g/kg EtOH (*p* = 0.0819). **F)** No significant impact was observed on event prominence in the first 5 minutes; *F*(1.172, 6.740) = 0.518; *p* > 0.05 (Greenhouse-Geisser corrected one-way repeated measures mixed-effects analysis). **G)** For saline, raster plots (top) are shown where each row represents an individual mouse, and each vertical line represents an identified astrocyte calcium event across the 30-minute open field session. Average astrocyte calcium events binned in one-minute bins (middle), and average distance traveled with the same binning (bottom) allow for visualization of concurrent astrocyte calcium and locomotor behavior. **H)** The same information shown in G, but when mice were given 2 g/kg EtOH. **I)** Alcohol inhibited astrocyte calcium events across the full 30-minute session, *F*(2.774, 15.95) = 4.938, *p* = 0.0144 (Greenhouse-Geisser corrected one-way repeated measures mixed-effects analysis). Tukey’s multiple comparison post-hoc tests indicated a significant difference between saline and 2 g/kg EtOH (*p* < 0.05). **J)** Event prominence was decreased when events from the full 30-minute session were considered, *F*(2.593, 14.91) = 5.230, *p* = 0.0139 (Greenhouse-Geisser corrected one-way repeated measures mixed-effects analysis). Tukey’s multiple comparison post-hoc tests indicated no significant differences between any two given doses. **K)** Overview of experimental design for open field tests created with cartoon mouse modified from SciDraw.io. **L)** Representative track plots in the first 5 minutes after mouse was given saline or **M)** 1 g/kg EtOH. **N)** Alcohol stimulation, as measured by distance travelled, peaked in the early part of the open field session during fiber photometry recordings; significant main effect of minute, *F*(1.738, 29.54) = 13.65, *p* = 0.0001, dose, *F*(1.000, 17.00) = 6.099, *p* = 0.0244, and dose x minute interaction, *F*(3.197, 47.32) = 9.170, *p* < 0.0001; Šídák’s multiple comparisons post-hoc tests indicated more movement at the 5 minute bin when mice were given 1 g/kg EtOH as compared to saline (*p* = 0.0076) (repeated measures mixed effects analysis). Note that this graph also includes behavioral data from mice recorded in the DMS (n = 18 total; DMS recording data shown in Figure S2). All error bars represent standard error of the mean (SEM).

**Figure 2.**
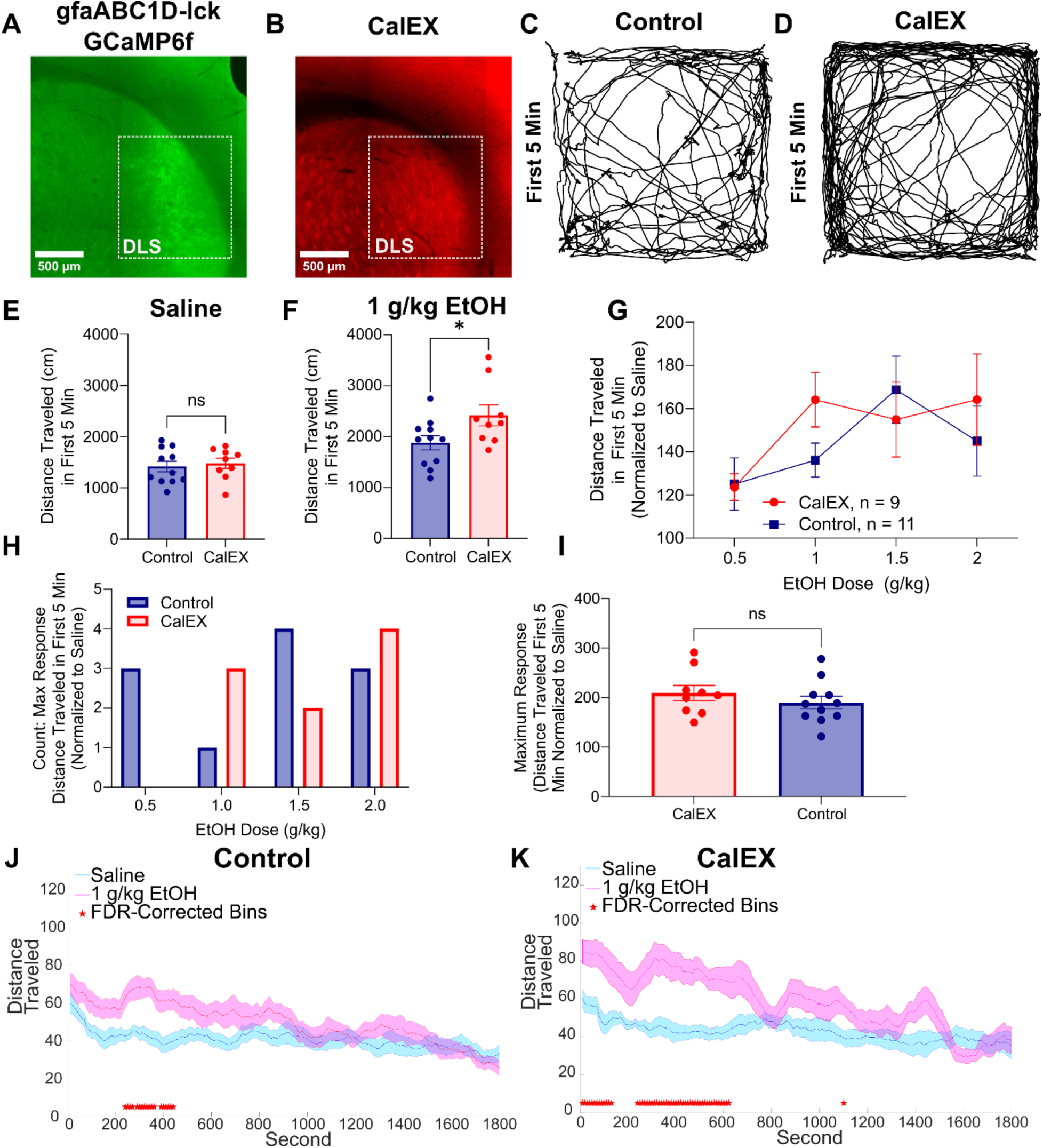
DLS astrocyte calcium activity inhibition increases the EtOH-induced stimulant response. **A)** Representative images of GCaMP6f-lck (control) and **B)** CalEX DLS expression. **C)** Representative track plots in the first 5 minutes when control or **D)** CalEX mice are given 1 g/kg EtOH. **E)** When given saline, viral groups moved the same in the first 5 minutes, *t*(18) =0.4144, *p* = 0.683, *d* = -0.188 (unpaired t-test). **F)** When given 1 g/kg EtOH, CalEX mice mounted a greater EtOH-induced stimulant response in the first 5 minutes, *t*(18) =2.228, *p* = 0.038, *d* = -0.985 (unpaired t-test). **G)** Across doses, this effect was most prominent at 1 g/kg EtOH, trend for a main effect of dose, *F*(3, 54) = 2.581, *p* = 0.0629, η^2^p = 0.125 (two-way repeated measures ANOVA). **H)** Mice displayed considerable variability in which EtOH dose produced the maximum EtOH-induced stimulant response (as determined using distance traveled in first 5 minute normalized to saline data); Fisher’s exact test comparing CalEX and control distributions: *p* = 0.2972. **I)** Maximum response regardless of EtOH dose did not differ between CalEX and control mice; *t*(18) = 0.9623, *p* = 0.3487 (unpaired t-test). **J)** Movement patterns under saline and 1 g/kg EtOH conditions reveal that control mice do display an EtOH-induced stimulant response; red stars indicate significant FDR-corrected t-tests for the 10-second time bins. **K)**. CalEX mice mounted a greater stimulant response that also lasted longer across the 30-minute open field session. All error bars represent standard error of the mean (SEM).

**Figure 3.**
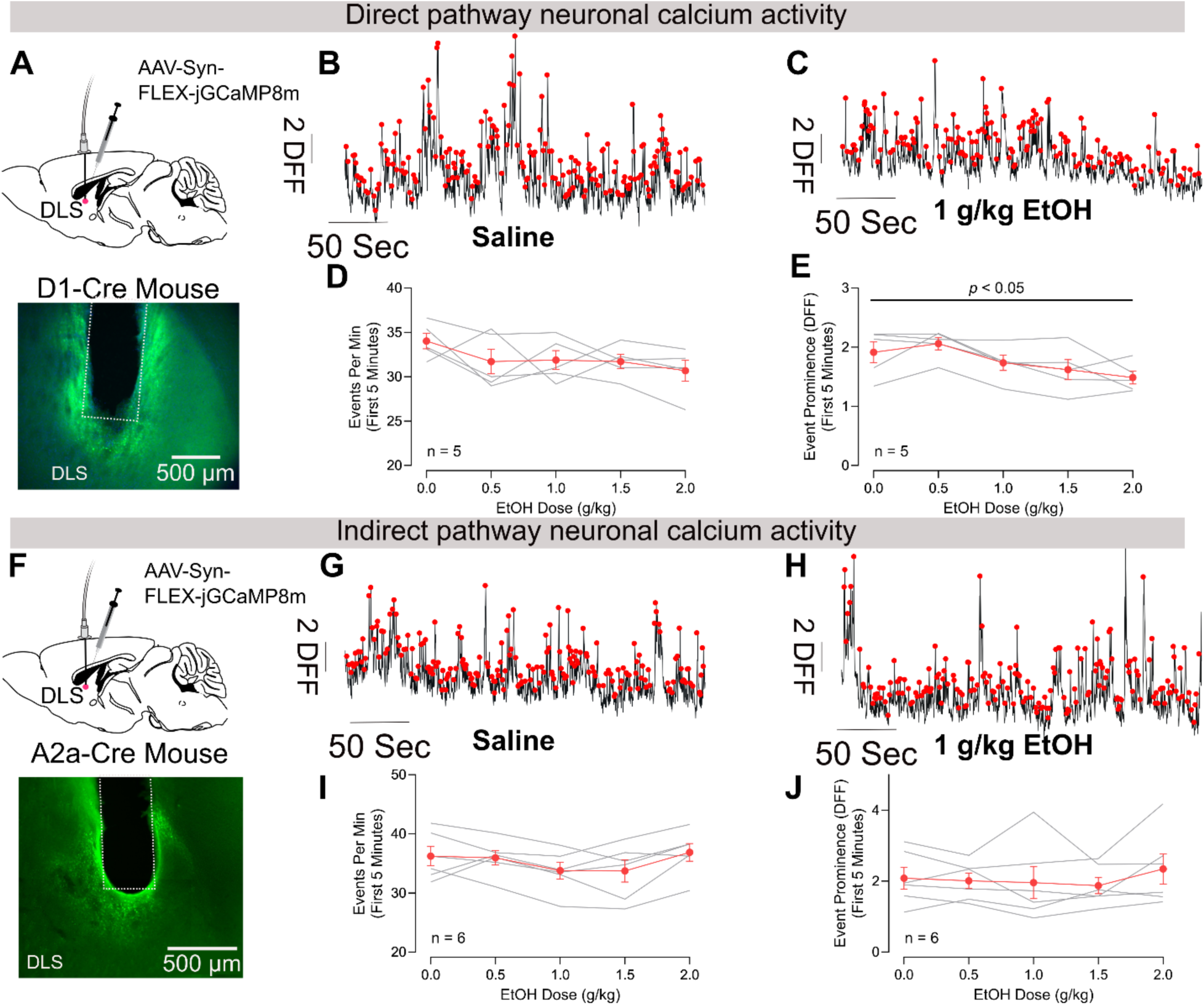
Acute alcohol exhibited minimal effects on striatal SPN activity. **A)** To record direct pathway neuron calcium activity, D1-Cre mice were given a stereotaxic injection of a Cre-dependent GCaMP virus with accompanying fiber optic implant for in-vivo fiber photometry calcium recordings (top); representative viral expression (bottom). **B)** Example direct pathway activity when mice were given saline or **C)** 1 g/kg EtOH. Red dots represent detected neuronal events. **D)** Acute alcohol did not have a significant impact on D1 neuron event rate; *F*(2.387, 9.549) = 1.700, *p* > 0.05, η^2^p = 0.298 (Greenhouse-Geisser corrected repeated measures one-way ANOVA); **E)** but did decrease event prominence at higher doses; *F*(1.795, 7.180) = 6.180, *p* = 0.029, η^2^p = 0.61 (Greenhouse-Geisser corrected repeated measures one-way ANOVA). **F)** To record indirect pathway neuron calcium activity, A2a-Cre mice were given a stereotaxic injection of a Cre-dependent GCaMP virus with accompanying fiber optic implant for in-vivo fiber photometry (top); representative viral expression (bottom). **G)** Example indirect pathway activity when mice were given saline or **H)** 1 g/kg EtOH. Red dots represent detected neuronal events. **I)** Acute alcohol did not have a significant impact on indirect pathway neuronal event rate; *F*(2.544, 12.72) = 3.01, *p* = 0.075, η^2^p = 0.376 (Greenhouse-Geisser corrected repeated measures one-way ANOVA); **J)** or prominence; *F*(2.161, 10.81) = 0.813, *p* = 0.478, η^2^p = 0.139 (Greenhouse-Geisser corrected repeated measures one-way ANOVA). All error bars represent standard error of the mean (SEM).

#### Open field video analysis

Open field video data were analyzed using DeepLabCut [42]. A model was trained to track six key points on the mouse: nose, right ear, left ear, body center, end of the tail, and base of the tail. Distance traveled was computed using the body center key point. Open field box corners were also tracked. Like analysis conducted by our lab previously [40], these points were used to calculate the length of each side of the box in pixels. X-Y pixel coordinates of the body center key point were converted to cm using the known cm length of each box side (40 cm). Points tracked with a < 99% probability were eliminated, and the remaining data were interpolated using the “interp1” MATLAB function. X-Y coordinates were LOESS filtered using the function “smooth” with a window size of 10 frames (0.5s). Coordinates were then used to calculate the frame-by-frame distance, which was summed to calculate the total distance traveled across various time bins.

### Effect of CalEX astrocyte expression on EtOH stimulation and other EtOH-related behaviors

#### CalEX: EtOH-induced stimulant response testing

To assess if DLS astrocyte calcium signaling plays a role in modulating the EtOH-induced stimulant response, we used CalEX to inhibit DLS astrocyte calcium signaling. CalEX is a calcium extrusion hPMCA pump, which works to constitutively extrude cytosolic calcium from the astrocyte soma [43]. This tool has been extensively validated [44–47], including in the striatum [43, 47]. In our first CalEX experiment, C57BL/6J mice were given bilateral stereotaxic DLS injections of gfaABC1D-GCaMP6f-lck as a control virus (n = 11; 6F, 5M) or CalEX (n = 9; 6F, 3M). Following recovery and time for viral expression (∼4 weeks), mice underwent EtOH-induced open field stimulant response testing as described above.

Using CalEX, we also assessed the role of DLS astrocytes in modulating the following additional EtOH-related behaviors: two-bottle choice home cage EtOH drinking, EtOH metabolism, drinking-in-the-dark (DID), aversion-resistant EtOH drinking, and loss of righting reflex (LORR). Please see Supplementary Note 1 for detailed methods and results of these additional behavioral assays.

### Neuromodulator release recordings during tail suspension test & open field locomotion

To assess how acute EtOH impacts striatal acetylcholine (ACh) and norepinephrine (NE) release, fiber photometry recordings were conducted as described above. C57BL/6J mice (n = 6, 5F, 1M) received a stereotaxic injection of AAV-hSyn-GRAB_ACh3.0 and fiber implant in the DLS. As a control for the GRAB-ACh, AAV-hSyn-ACh4.3_mut (n = 2M mice targeting DLS) was used as a null sensor. To measure NE release, C57BL6J (n = 4F) received a stereotaxic injection of rAAV-hSyn-GRAB-NE2m(3.1)-WPRE-pA and fiber in the DLS. Another group of C57BL6J mice (n = 3; 1F, 2M) targeting the BNST [AP = +0.14, ML = ± 0.88, DV = -4.24] [48]) was included as a control brain region which receives substantial noradrenergic input [49].

In addition to open field EtOH-induced stimulant response testing, these mice also underwent a tail suspension test with concurrent fiber recordings the week after stimulant response testing. This test was performed as 5 trials that lasted 60 seconds each with a minimum of 60 seconds in between each trial [50]. The tail suspension test occurred in the open field because this apparatus was already outfitted for fiber photometry recordings.

### Chemogenetic manipulation of cholinergic interneurons (CINs)

#### Impact on ACh-release

ChAT-Cre mice (n = 5; 2F, 3M) received an injection of AAV-hSyn-GRAB_ACh3.0 in the cortex (AP: +2.1, ML: +0.4, DV: - 2.3). One week later, mice received a stereotaxic injection of a Cre-dependent Gq-DREADD virus (AAV-hSyn-DIO-hM3D(Gq)-mCherry) along with an optical fiber in the DLS. By expressing the GRAB-ACh sensor in cortical axons projecting to the the striatum instead of injecting it locally in striatum, we would overcame known issues co-expressing two viruses. This facilitated the ability to activate CINs with simultaneous measurement of ACh release. These mice received a pre-treatment with CNO (1 mg/kg) or saline 50-minutes before a 10-minute baseline fiber recording, following which mice were given an additional i.p. injection of saline. CNO was prepared as a 0.1 mg/mL solution in 0.9% sterile saline.

#### Impact on astrocyte calcium activity

A separate group of ChAT-Cre mice (n = 10, 1F, 9M) received a stereotaxic co-injection of a Cre-dependent Gq-DREADD virus (AAV-hSyn-DIO-hM3D(Gq)-mCherry) mixed 1:1 with an astrocyte-specific GCaMP virus (AAV-gfaABC1D-cyto-GCaMP6F) along with an optical fiber in the DLS. This facilitated the ability to activate CINs with simultaneous measurement of DLS astrocyte calcium. Another group of ChAT-Cre mice (n = 4; 2F, 2M) served as the viral control, receiving a Cre-dependent mCherry (AAV-hSyn-DIO-mCherry) instead of the DREADD along with the astrocytic GCaMP and DLS fiber. All mice underwent EtOH-induced stimulant response testing with a 30-minute DCZ (0.2 mg/kg, [51]) or saline pre-treatment and 10-minute baseline fiber recording preceding the EtOH or saline injection (1 g/kg EtOH). Then, half of the DREADD mice underwent the same protocol, but now testing for 2 g/kg EtOH. These mice then underwent subsequent LORR testing as described above. A subset of the mice underwent head-fixed motorized wheel testing, as described by our lab previously [40]. Mice were first acclimated to a head-fixed voluntary running wheel for 3 days. Then, on a given testing day, they underwent 25 motorized locomotion trials (10s duration) with at least 50 seconds of rest in between. On these testing days, the mice received an i.p. injection of DCZ (0.2mg/kg) or saline 30-minutes prior to the start of the session and a subsequent i.p. injection of saline immediately before testing began.

### Perfusion, immunofluorescent staining, and histological imaging

At the conclusion of all experiments, mice were transcardially perfused with heparinized 1X phosphate buffered saline (PBS) followed by 4% paraformaldehyde (PFA). Brains were extracted and stored in 4% PFA in a 4°C refrigerator overnight and then transferred to 1X PBS the following day where they remained in the refrigerator until they were sliced using a vibrating microtome (Compresstome, Precisionary). Brain slices containing the targeted brain regions were assessed for fiber optic and viral placement, or viral placement only, and were sliced at 100 µM. Brains that were used for immunofluorescent staining (described below) were sliced at 60 µM. All brain slices were imaged using a Widefield Leica DMI8 epifluorescence microscope with automated tile stitching. Mice that were identified to have misplaced viral and/or optical fiber placements were excluded from all analyses and are not reflected the reported sample sizes throughout the manuscript.

For mice that received GRAB sensor (or control) stereotaxic injections, brains were counterstained with GFP to boost the signal. All steps were completed with slices shaking at room temperature unless otherwise noted. Brains were sliced at 60 µM and then washed 3 times in 1X phosphate-buffered saline (PBS) for 15 minutes each wash. Then, slices were incubated in 50% methanol in 1X PBS for 30 minutes. Slices were then incubated in 1% hydrogen peroxide in 1X PBS for 15 minutes then washed 3 times in 1X PBS for 15 minutes each wash. Slices were then washed with 1X PBS + 0.4% Triton-X for 30 minutes and then blocked in 10% normal goat serum (NGS) in 1X PBS + 0.2% Triton-X for 90 minutes. Following blocking, slices were incubated in the primary antibody (1:500 of GFP Polyclonal Antibody from ThermoFisher, #A11122; in NGS +0.02% Triton-X) overnight at 4°C while shaking. The next day, brains were washed 4 times in 1X PBS for 10 minutes each wash. Then, they were incubated in secondary antibody for 2 hours (Goat anti-Rabbit IgG (H+L) Highly Cross-Adsorbed Secondary Antibody, Alexa Fluor™ 647, ThermoFisher, # A-21245; 1:1000 dilution in 1X PBS). Slices were then washed 3 times in 1X PBS for 10 minutes each wash and then mounted on glass slides and dried in the dark overnight. Once dry, slides were cover slipped and imaged using a Widefield Leica DMI8 epifluorescence microscope.

### Analyses and consideration of sex effects

For slice 2P imaging and fiber photometry, events were detected as described above. Event rate and amplitude or prominence was analyzed across EtOH conditions using one-way repeated measures ANOVAs (or mixed effects analysis in the case of missing data; e.g., a documented missed EtOH injection) using GraphPad Prism 11.0.0. For consideration of the EtOH-induced stimulant response, locomotor data were processed using DeepLabCut as described above and analyzed across EtOH and viral groups using one-way or two-way repeated measures ANOVAs as applicable. T-tests were utilized for some comparisons (i.e., LORR dependent variables comparing CalEX vs. viral control groups). Throughout the manuscript, post-hoc analyses were conducted when warranted and Greenhouse-Geisser corrections were applied to analyses where appropriate.

Where possible, sex effects were considered and reported in the results section if found. Some fiber photometry experiments have a low sample size between sexes, which precluded analysis by sex (see Supplementary Table 2 for sample sizes for each experiment). If there were no interaction effects with sex, results are shown collapsed on sex and null sex-based analyses are reported in Supplementary Table 3.

## Results

### Acute EtOH inhibits in vivo dorsolateral striatal astrocyte calcium signaling

We used two-photon calcium imaging to begin assessing the impact of acute EtOH on striatal astrocyte calcium activity. We virally expressed GCaMP6f in the DLS under the control of an astrocyte-specific promoter and prepared acute ex vivo brain slices for imaging (Fig. S1A). Consistent with previous work, astrocytes showed robust intracellular calcium dynamics in the absence of any experimental stimuli under baseline conditions (Fig. S1B, C). We bath applied 20 and 50 mM EtOH followed by a washout to determine how EtOH impacts this spontaneous calcium activity. EtOH had no significant effect on astrocyte activity metrics quantified by detecting individual calcium transients (red dots in Fig. S1C). Event frequency and amplitude were similar across baseline, EtOH, and washout conditions (Figure S1D-G). These findings show that EtOH has minimal effects on astrocyte calcium activity recorded in a reduced preparation lacking the full complement of DLS synaptic inputs, neuronal activity and neuromodulation dynamics.

In vivo fiber photometry was then used to capture population-level bulk activity dynamics in the intact striatal circuit. This strategy also allowed us to assess EtOH -induced stimulation in parallel with bulk activity changes. We first report how acute EtOH changes in vivo astrocyte activity followed by analysis of how these changes relate to EtOH-induced stimulation. We virally expressed in DLS astrocytes the calcium sensor gfaABC1D-GCaMP6f and implanted an optic fiber above the viral injection site (Figure 1A, B). Mice were given intraperitoneal (i.p.) injections of saline (control) or EtOH (0.5, 1, 1.5, and 2 g/kg) in separate daily sessions and placed immediately in an open field box with an overhead video camera (Figure 1C). We restricted our initial activity analysis to the first 5 mins after the i.p. injection since previous work shows EtOH stimulation effects in this same time window [25, 52]. Astrocytes showed robust calcium dynamics under baseline saline conditions in freely moving animals (Figure 1D), similar to our recent observations in head-fixed mice [40]. Examining astrocyte calcium activity traces recorded across doses suggested suppression of astrocyte activity with EtOH (Figure 1D). We identified individual calcium events and quantified event frequency and prominence (amplitude relative to surrounding events) as a measure of astrocyte activity. In contrast to our observations in brain slices, EtOH acutely inhibited in vivo striatal astrocyte calcium activity by reducing event frequency (Figure 1E), but event prominence was unaffected (Figure 1F).

Given that our observed astrocyte activity inhibition and the previously reported stimulant response both occur during the first 5 mins after EtOH injection, we examined the temporal relationship between these EtOH-induced effects in our data. We used DeepLabCut to track mouse location in fiber photometry open field videos. We plotted identified astrocyte calcium events as raster plots and quantified average events per minute across the full 30-minute open-field session. When these data were plotted alongside distance traveled a temporal relationship between these two variables emerged, such that when astrocyte calcium activity was inhibited by EtOH, the mice tended to show acute EtOH stimulation (i.e., they moved more than when given saline) (Figure 1G, H). Across the full 30-minute session, astrocyte calcium events and prominence were inhibited by EtOH (Figure 1I, J). Notably, performing this same protocol in the DMS produced similar results (Figure S2), suggesting that the EtOH-induced astrocyte calcium signaling inhibition observed in vivo is robust across the striatum and not DLS-specific. We focused our further studies in the DLS given the well-established role of this striatal subregion in EtOH stimulation. Taken together, these results show that EtOH reduces astrocyte activity in the intact striatal circuit.

### EtOH induces behavioral stimulation in parallel with astrocyte activity inhibition

EtOH exposure recruits a stimulant response that manifests as increased locomotion in rodents [25, 53]. We further examined the stimulant response under our fiber photometry recording conditions. Because the DLS and DMS recording cohorts are the same mouse strain (C57BL/6J) and the behavioral testing was conducted identically, we pooled the behavioral data from both groups to increase the sample size. Indeed, distance traveled in the first 5 minutes normalized to saline locomotion across all tested doses (Figure S3A) indicated no significant main [*F*(1,16) = 0.423, *p* = 0.52] or interaction effect with brain region [*F*(1.848, 26.49) = 0.269, *p* = 0.75]; Greenhouse-Geisser corrected two-way (brain region x dose) mixed effects analysis. Locomotion was quantified as the open field distance traveled as a function of time from injection (see Methods). Consistent with previous findings in untethered animals, there was higher locomotion in response to an i.p. injection of 1g/kg EtOH relative to saline injections (Figure 1K - N). This stimulation effect peaked in the first 5 mins after EtOH injection, during the same time when astrocyte activity inhibition was observed.

EtOH-induced stimulation, quantified as the distance traveled in first 5 minutes normalized to saline locomotion, was highly variable across mice (Figure S3A), which is consistent with previous reports in both humans [14, 19] and rodents [25]. Due to this variability, there was no significant difference in EtOH-induced stimulant response across doses (Figure S3A). Relatedly, we found that there was no single dose where mice as a group had the maximal or minimal EtOH-induced stimulant response (Figure S3B), further highlighting variability in individual EtOH responses. Taken together, our behavioral analyses show that tethered mice undergoing fiber photometry recordings show a robust EtOH stimulation response with considerable interindividual differences. Given the important role of astrocyte calcium signaling in multiple homeostatic and neuromodulatory processes [54], these results raise the possibility that EtOH’s behavioral and neurophysiological effects are modulated by its suppression of astrocyte activity. We addressed this possibility in our next series of experiments.

### DLS astrocyte calcium inhibition facilitates the EtOH-induced stimulant response

To establish how astrocyte calcium activity contributes to EtOH stimulation, we virally overexpressed the molecular genetic tool CalEX selectively in this cell type. This virus encodes a human plasma membrane Ca^2+^-ATPase that decreases astrocyte calcium activity [43] and thus mimics the suppressive effect of EtOH on astrocytes in a constitutive manner. C57BL/6J mice were given bilateral stereotaxic DLS injections of gfaABC1D-CalEX or gfaABC1D-GCaMP6f-lck as a fluorophore control (Figure 2A, B). These experiments were performed similarly as above, except no mouse tethering for calcium recording was required. Inspecting 2D locomotion traces of individual mice suggested that EtOH induces a greater stimulant response in CalEX expressing animals as compared to viral controls (Figure 2C, D). Viral groups did not differ in baseline locomotion as there was no difference in total distance traveled in the first 5 mins after saline injections (Figure 2E). However, when given 1 g/kg EtOH, CalEX expressing mice showed greater EtOH-induced stimulation (Figure 2F). When considering the stimulant response across the range of tested doses, CalEX expression seemed to shift peak stimulation to a lower dose compared to controls (Figure 2G). To address if this shift is due to changes in EtOH sensitivity, we quantified for each viral group the number of mice showing their maximal stimulant response at each dose. Mice in both viral groups showed considerable variability in the dose which produced a maximally stimulating response, and there was no difference in the distribution of maximally stimulating dose between groups (Figure 2H). This suggests that the observed enhancement in stimulation is unlikely due to changes in EtOH sensitivity. In agreement, comparing the maximal stimulant response observed for each animal regardless of the dose at which it occurred showed no difference between control and CalEX mice (Figure 2I). Taken together, these findings suggest that attenuating astrocyte calcium activity with CalEX potentiates the stimulant response at submaximal doses without affecting the dose at which the maximal stimulant response is produced.

Finally, we tested the impact of astrocyte CalEX expression on the duration of the stimulant response. We quantified locomotion across the full recorded 30 mins for saline and 1g/kg EtOH sessions. Control mice displayed the expected EtOH-induced stimulant response (Figure 2J), however, in CalEX-expressing mice, EtOH stimulation was more pronounced and lasted for longer (Figure 2K). Together, these results indicate that DLS astrocyte calcium activity modulates the intensity and duration of the EtOH-induced stimulant response.

Given the above results, in 2 new cohorts of mice we tested whether striatal astrocyte CalEX expression impacts other EtOH-related behaviors (see Supplementary Note 1 for full description). In the first cohort, we tested mice in home-cage intermittent 2-bottle choice (2BC) drinking using automated Promethion (Sable) behavioral monitoring cages (Figure S4). In the next cohort, we assayed whether CalEX and control viral groups differ in EtOH metabolism, drinking-in-the-dark (DID) home cage drinking, aversion-resistant drinking, and loss of righting reflex (Figure S5). Importantly, there were no differences between viral groups in EtOH metabolism (Figure S5D), suggesting that our findings related to EtOH-induced stimulation are not driven by differences in metabolic rate. Overall, these experiments showed minimal effects of CalEX astrocyte expression on EtOH sedation and EtOH consumption behaviors, suggesting that DLS astrocyte calcium signaling specifically modulates EtOH stimulation.

### EtOH exerts minor effects on bulk striatal projection neuron activity

Our ex vivo two photon calcium imaging experiments showed minimal effects of EtOH on spontaneous astrocyte calcium activity, yet we observed robust inhibition in our fiber photometry experiments. In vivo astrocyte calcium activity is strongly dependent on local neuronal activity and neuromodulatory inputs [54, 55], which are compromised in the ex vivo brain slice system. We reasoned that the discrepancy between our ex vivo and in vivo findings suggest the involvement of EtOH-sensitive circuit-level processes in the observed inhibition of astrocyte activity. Striatal astrocytes express a variety of neurotransmitter receptors [54, 56, 57], including GPCR-coupled GABAB receptors which enable these cells to respond to GABA release from local SPNs with intracellular calcium activity [58]. Therefore, the EtOH-induced astrocyte activity inhibition we observed could reflect changes in SPN activity. To consider this possibility, transgenic Drd1-Cre and A2a-Cre mice were given a stereotaxic injection of a Cre-dependent GCaMP8m calcium sensor in the DLS to label direct and indirect pathway SPNs, respectively (Figure 3). We performed fiber photometry open field recordings to assess the effect of EtOH on SPN activity. There was no effect of EtOH on the event rate of direct pathway neuron activity (Figure 3A-D) but event prominence was mildly reduced (Figure 3E). In contrast, we could not detect an effect of EtOH on indirect pathway neuron activity (Figure 3F-J). Hence, EtOH has minimal effects on DLS SPN activity, which is unlikely to explain the observed astrocyte suppression.

### Acute EtOH inhibits DLS acetylcholine release

Given the minimal effect of EtOH on SPN activity, we next tested the involvement of neuromodulatory inputs. EtOH inhibits cerebellar and cortical astrocyte calcium activity via inhibition of norepinephrine (NE) release [34]. Though the DLS receives only sparse NE innervation [59], we considered if the mechanism of EtOH-induced astrocyte calcium signaling could be the same as previously described. We virally expressed the genetically encoded GRAB-NE2m sensor [60] and used fiber photometry to measure NE release. Given potential concerns with crosstalk of GPCR based NE sensors [59], we first validated our use of the GRAB-NE sensor in a control brain region known to receive strong NE innervation (bed nucleus of stria terminalis, BNST) [49]. We virally expressed GRAB-NE and implanted a fiber in the BNST (Figure S6A, B) or in the DLS (Figure S6C, D). Tail suspension leads to high NE release [50]. Therefore, we determined NE release dynamics in the BNST and the DLS during tail suspension (Figure S6E). As expected, recordings from the BNST displayed robust NE release during tail suspension (Figure S6F). In contrast, we could not detect substantial NE release in the DLS in response to the same manipulation. (Figure S6G). Accordingly, direct comparison of recordings in the two brain regions showed significantly higher NE release in the BNST as compared to the DLS. (Figure S6H). Overall, this series of experiments establish a minimal NE tone in the DLS. Hence, the observed inhibition of astrocyte calcium activity likely involves a mechanism specific to the neuromodulatory landscape of the striatal circuit.

Striatal activity is under strong neuromodulatory control from local cholinergic interneurons (CINs) that tonically release acetylcholine (ACh) across the striatum [61]. Astrocytes in the dorsal striatum are found in close proximity to CINs [62], which may represent a structural basis for functional communication. Recent work shows acute EtOH inhibits ACh release in the dorsomedial striatum [63], where EtOH also suppressed astrocyte calcium activity (Figure 4). We tested the possibility that astrocyte inhibition by EtOH involves cholinergic signaling. We began by assessing how acute EtOH impacts in vivo ACh release in the DLS, which previous work did not address. Mice were given a stereotaxic DLS injection of GRAB-ACh3.0 (Figure 4A, B) or the null (ACh4.3_mut) sensor as control (Figure 4C) along with an optical fiber implant for fiber photometry recordings. Mice were placed in an open field for 10 mins to record baseline GRAB-ACh activity, after which they received i.p. injections of saline or EtOH. Acute injected EtOH led to a pronounced decrease in DLS ACh release across the range of tested doses (Figures 4D-F). This phenomenon was not observed in mice expressing the control null sensor (Figure 4G), indicating that EtOH exposure is not associated with non-specific changes in the fiber photometry fluorescence signal. As an additional fluorophore control, we tested mice expressing GRAB-NE in the DLS with saline and EtOH injections. Acute EtOH injections did not produce changes in the GRAB-NE signal (Figure S6 I-K), further validating the described astrocyte and ACh release EtOH effects.

**Figure 4.**
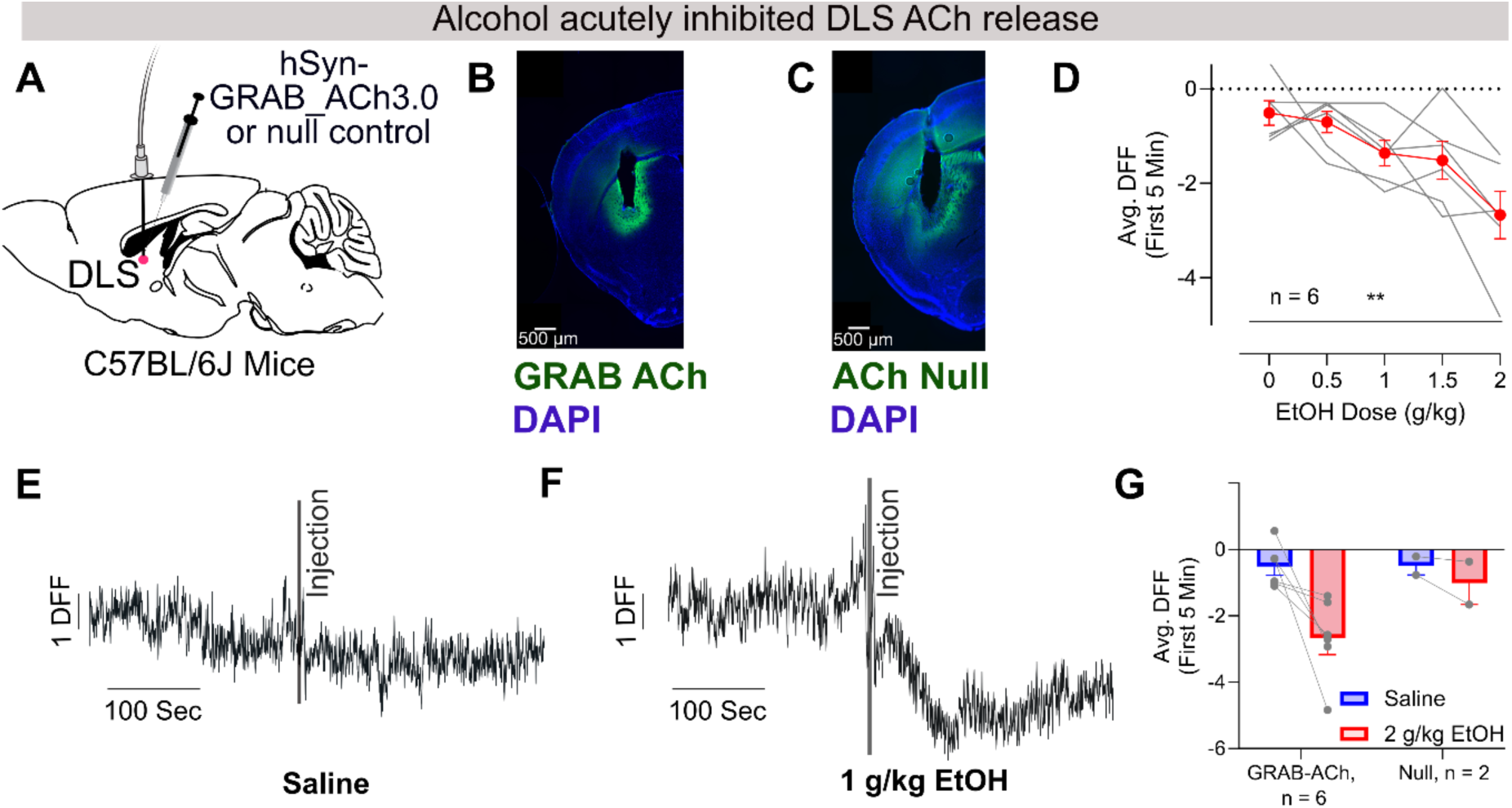
Acute alcohol inhibited DLS acetylcholine release. **A)** C57BL/6J mice received a stereotaxic viral injection of GRAB-ACh3.0 or null control sensor (AAV-hSyn-ACh4.3_mut) and an optical fiber for recording. **B)** Representative image of DLS GRAB-ACh and fiber placement. **C)** Representative image showing null control sensor expression. **D)** In the first 5 minutes, EtOH inhibited ACh release as measured by average DFF, *F*(2.175, 10.88) = 7.488, *p* = 0.0081 (Greenhouse-Geisser corrected repeated-measures one-way ANOVA). **E)** Example ACh DFF trace when a mouse received saline, or **F)** 1 g/kg EtOH. **G)** Mice expressing a control null ACh sensor in the DLS did not show as pronounced of a decrease in avg. DFF in the first 5 minutes from saline to the highest dose of EtOH tested (2 g/kg); GRAB-ACh: *t*(5) = 3.216, *p* = 0.0235; Null = *t*(1) = 1.402, *p* = 0.3944 (paired t-tests within virus). All error bars represent standard error of the mean (SEM).

### Chemogenetic stimulation of DLS cholinergic interneurons has no effect on astrocyte calcium activity

Our results so far show that EtOH inhibits both DLS astrocyte calcium activity and ACh release, raising the possibility that astrocyte activity suppression is driven by the concomitant decrease in extracellular ACh. Such a mechanism implies a facilitatory effect of ACh release on astrocyte calcium activity under basal conditions. We tested this prediction by first examining how Gq-DREADD stimulation of CINs affects ACh release. ChAT-Cre mice received a stereotaxic injection of Cre-dependent hM3Dq-mCherry in the DLS. We virally delivered GRAB-ACh to the cortex so it is expressed in corticostriatal axons (Figure S7A, B). This strategy allowed us to measure ACh release by monitoring fluorescence changes in axonally expressed GRAB-ACh while avoiding potential issues with viral co-expression within the striatum itself (Figure S7B). Mice were given an i.p. injection of the DREADD agonist clozapine-n-oxide (CNO, 1mg/kg) and 50 mins later put in an open field box for recordings. Gq-DREADD stimulation of CINs with CNO did not change the frequency or prominence of ACh release events as compared to the control saline injection (frequency: *t*(4) = 0.5928, *p* = 0.5852; prominence: *t*(4) = 0.3572, *p* = 0.7390). Surprisingly, we observed that a subsequent i.p. injection, which involved scruffing the animal, increased the ACh release event rate in mice previously injected with CNO compared to saline (Figure S7G). These findings suggest that Gq-GPCR based stimulation of CINs does not appreciably increase ACh release under basal conditions but sensitizes ACh release to subsequent stimuli like scruffing.

Given that we could increase ACh release with DREADD stimulation of CINs, we sought to determine how this manipulation affects DLS astrocyte calcium activity. In a new group of ChAT-Cre mice, we delivered into the DLS a viral mixture to co-express astrocytic GCaMP and Cre-dependent Gq-DREADD-mCherry (or AAV-hSyn-DIO-mCherry as control) in CINs, and implanted an optical fiber for photometry recordings (Figure 5A, B). Mice were given an i.p. injection of the DREADD agonist DCZ (0.2 mg/kg) or saline as control and 50 mins later placed into an open field box for fiber photometry recordings. After recording baseline astrocyte calcium activity for 10 mins, mice were given an i.p. injection of saline (Figure 5) or EtOH (Figure 6) and placed back into the box to continue recording. We analyzed activity after the second i.p. injection since this is when we observed increased ACh release in the previous experiment (Figure S7). There was no difference in astrocyte event frequency (Figure 5C – E) or prominence when comparing the same mice pretreated with DCZ or saline (prominence data not shown; sal-sal, mean prominence = 1.312, std. dev = 0.3493; DCZ-sal, mean prom. = 1.356, std. dev. = 0.3694; *t*(9) = 0.4946, *p* = 0.6327, paired *t*-test). We also did not detect any changes in astrocyte activity in control mice expressing mCherry in CINs (Figure 5E). These results suggest that Gq-DREADD stimulation of CINs has minimal effects on astrocyte calcium activity during open field exploration.

**Figure 5.**
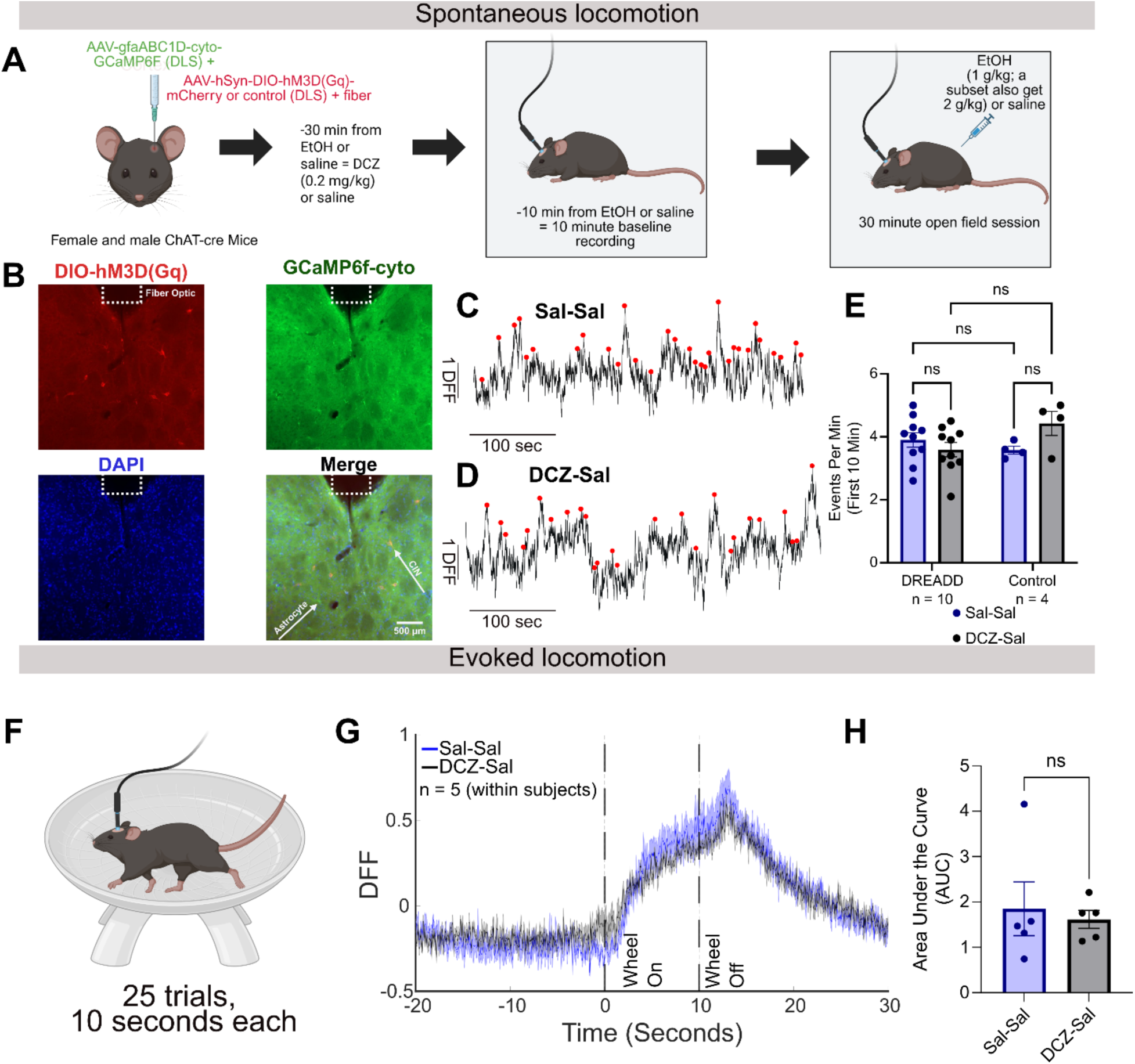
Stimulation of cholinergic interneurons using DREADDs did not impact astrocyte calcium activity during spontaneous or evoked locomotion. **A)** ChAT-Cre mice received a stereotaxic co-injection of gfaABC1D-cyto-GCaMP6f with either a Cre-dependent Gq-DREADD (AAV-hSyn-DIO-hM3D(Gq)-mCherry) or control virus (AAV-hSyn-DIO-mCherry) in the DLS. An optical fiber was placed above the injection site to measure astrocytic calcium activity. On a given day, mice received DCZ to activate cholinergic interneurons (CINs) (or received saline as a drug control) 30 minutes prior to 1 g/kg EtOH (or saline). Ten minutes prior to 1 g/kg EtOH (or saline), mice received a 10-minute baseline recording in the open field. Mice then received their EtOH or saline injection, and astrocytic calcium and overhead video were subsequently recorded for 30-minutes in the open field. A subset of these mice were also tested the following week using this same protocol, but now instead with 2 g/kg EtOH. Note that this figure is only displaying data from the saline treatment, which is pulled from this larger experiment. The EtOH data is featured in the next figure. **B)** Representative image showing fiber placement, expression of DREADD (red) and GCaMP6f-cyto (green), DAPI (blue), and these three merged. In the merged image, example CINs and astrocytes are labeled. **C)** Representative astrocyte calcium traces when mice received saline-saline or **D)** DCZ-saline. **E)** Stimulation of CINs via DCZ did not impact astrocytic calcium activity event rate in the first 10 minutes of the open field session in DREADD or viral control mice; a significant virus x treatment interaction was found, *F*(1, 12) = 6.073, *p* = 0.0298, η^2^p = 0.336 (two-way ANOVA), but post-hoc comparisons revealed no significant differences between and across conditions. **F)** To assess if CIN stimulation could modulate astrocyte calcium activity during evoked locomotion, a subset of mice (mice that did not undergo 2 g/kg EtOH testing) were head-fixed and tested using a motorized wheel, which prompted 25 10-second-long movement bouts. **G)** Percent DFF of astrocyte calcium activity during saline-saline vs. DCZ-saline testing indicated that CIN stimulation had no prominent impact on astrocytic activity. **H)** Quantification of area under the curve (AUC) from wheel on to wheel off (time 0 to 10 seconds on graph G) indicated no significant difference during CIN stimulation (DCZ-saline); *t*(4) = 0.322, *p* = 0.7635 (paired t-test). All error bars represent standard error of the mean (SEM). Panels A and F were created using BioRender.

**Figure 6.**
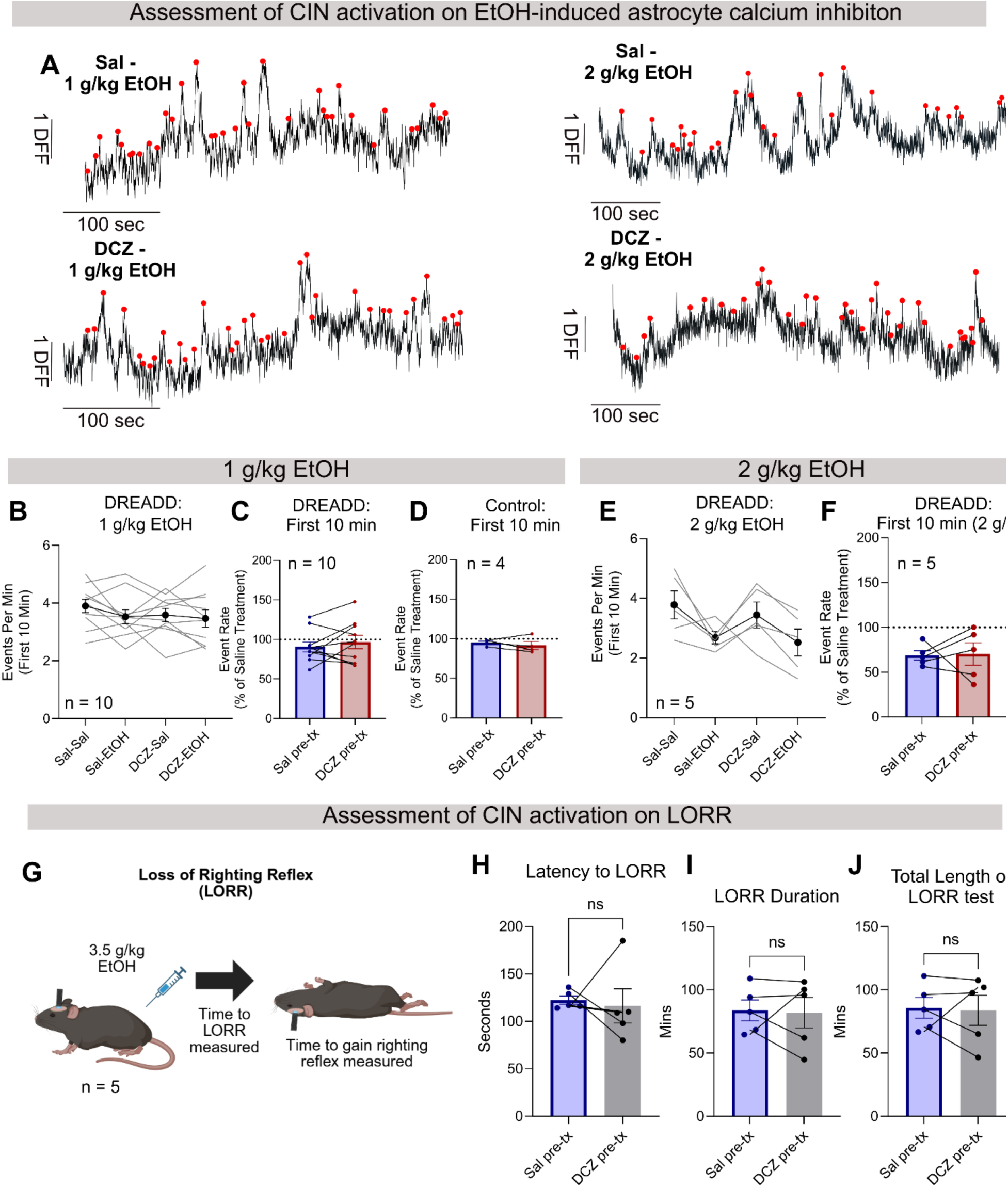
CIN stimulation does not reduce alcohol-induced astrocyte calcium activity inhibition. **A)** Astrocyte calcium traces from a representative mouse when receiving saline-EtOH (1 g/kg), saline-EtOH (2 g/kg), DCZ-EtOH (1 g/kg), and DCZ-EtOH (2 g/kg). **B)** In the first 10 minutes, no significant difference in event rate was noted across treatment groups, *F*(2.698, 24.28) = 1.186, *p* = 0.3329, η^2^p = 0.116 (Greenhouse-Geisser corrected repeated measures one-way ANOVA). **C)** Event rate in the first 10 minutes was normalized to within-treatment saline (i.e., saline-EtOH was normalized to saline-saline and DCZ-EtOH was normalized to DCZ-saline) to facilitate comparison across pre-treatment conditions. This analysis indicated that there was no significant difference in normalized event rate between the pre-treatment groups for mice expressing the active DREADD, *t*(9) = 0.8836, *p* = 0.3999 (paired t-test). **D)** Null results were also observed in control mice expressing only mCherry, *t*(3) = 0.8376, *p* = 0.4638 (paired t-test). **E)** In the subset of mice tested for 2 g/kg EtOH, we also observed no significant difference in event rate across treatment combinations in the first 10 minutes, *F*(1.956, 7.823) = 3.380, *p* = 0.0881, η^2^p = 0.2279 (Greenhouse-Geisser corrected repeated measures one-way ANOVA). **F)** No significant difference was observed between pre-treatment conditions when normalized to within-treatment saline, *t*(4) = 0.1008, *p* = 0.9245 (paired t-test). G) **J)** The subset of mice that underwent 2 g/kg EtOH testing subsequently underwent a LORR assessment to measure the impact of CIN stimulation on EtOH-induced sedation. **H**) No differences were observed in latency to LORR between saline and DCZ pre-treatments, *t*(4) = 0.2860, *p* = 0.7891 (paired t-test). **I)** No differences were observed in LORR duration, *t*(4) = 0.1810, *p* = 0.8651 (paired t-test). **J)** No differences were observed in total length of LORR test, *t*(4) = 0.1915, *p* = 0.8575 (paired t-test). All error bars represent standard error of the mean (SEM). Note: these are the same mice from which data are shown in Figure 5, now focusing on EtOH treatment.

Previous work from our lab [40] and others [64–67] shows robust recruitment of astrocyte calcium responses during locomotion and movement more generally. We did not find any changes in astrocyte calcium activity as a result of CIN stimulation during spontaneous locomotion in the open field. Behavior in the open field is dominated by diverse head and other body movements, which makes it challenging to specifically isolate the effect of CIN stimulation on locomotion-related astrocyte responses. We trained a subset of the same mice on a head-fixed motorized wheel paradigm (see Methods) and recorded astrocyte calcium signals via fiber photometry (Figure 5F). Mice were pretreated with DCZ or saline 30 mins before a subsequent i.p. injection of saline. Motorized locomotion elicited robust calcium responses that were similar with either DCZ or saline pre-treatment (Figure 5G – H). Together, these experiments show that under basal conditions, CIN stimulation has no measurable effects on astrocyte activity that are detectable by bulk fiber photometry.

CINs are tonically active, leading to a high ACh release rate in the striatum. One explanation for not observing astrocyte effects of increasing ACh release is that under basal conditions astrocyte cholinergic signaling is saturated. In this scenario, EtOH-induced suppression of ACh release could lead to astrocyte activity suppression. We reasoned that if this were the case, then augmenting ACh release with chemogenetic CIN stimulation should mitigate EtOH-induced changes in astrocyte activity. The same mice as described in Figure 5 were pretreated with either DCZ or saline and then subsequently injected with 1g/kg and, in a subset of mice, 2g/kg EtOH. We performed fiber photometry and compared astrocyte calcium activity across the tested conditions (Figure 6A). At 1 g/kg EtOH, we did not observe any significant differences across conditions (Figure 6B, C). This was also true in our viral control mice (Figure 6D). At 2 g/kg EtOH, there was more robust astrocyte calcium activity inhibition in the saline-EtOH group, however, CIN stimulation via DCZ pretreatment did not modulate this effect (Figure 6E, F).

Next, we considered the behavioral effects of CIN stimulation. We note that this experiment was designed primarily to address physiological effects of CIN manipulations on astrocytes and hence employed unilateral (and not bilateral) manipulations. With this caveat in mind, we determined how CIN stimulation affects EtOH stimulation. Examining locomotion behavior after i.p. injections of saline and 1g/kg EtOH suggested differences between the DCZ and saline pretreatment conditions (Figure S8A-C). We quantified EtOH stimulation as the locomotion observed with EtOH normalized by saline for each pretreatment condition. 1g/kg produced a variable stimulant response as before (Figure S8D). There was a trend for lower EtOH stimulation in the DCZ pretreatment condition as compared to saline (Figure S8D). Given these results, we also evaluated the impact of CIN stimulation on EtOH-induced sedation using the LORR test. We did not observe any impact of CIN stimulation on the tested LORR dependent variables (Figure 6G – J). The observed trend for reduced EtOH stimulation with CIN neuron activation is broadly consistent with previous pharmacological data showing potentiation of EtOH stimulation with muscarinic antagonism [68].

## Discussion

We found that acutely injected EtOH produced an expected locomotor stimulant response, which corresponded with an EtOH-induced inhibition of DLS astrocyte calcium signaling (Figure 1). Over-expression of CalEX in DLS astrocytes (which inhibits astrocytic calcium signaling [43] and thus mimic’s alcohol’s suppressive effect on these cells), resulted in a greater EtOH-induced stimulant response to 1 g/kg EtOH as compared to mice expressing a control virus (Figure 2). This finding suggests that DLS astrocytes play an important role in regulating EtOH-induced stimulation. Astrocyte CalEX expression did not change other alcohol-related behaviors including 2-bottle choice (2BC) home-cage drinking, drinking-in-the-dark (DID; a measure of binge alcohol drinking), aversion-resistant drinking, or loss of righting reflex, overall suggesting a somewhat specific role for DLS astrocytes in modulating EtOH-induced stimulation (Figures S4, S5).

These findings on DLS astrocytes are broadly consistent with known roles of the DLS itself in the development of alcohol drinking and alcohol use disorder. In addition to an established role in facilitating alcohol-induced stimulation [30], the DLS plays a key role in habitual alcohol drinking and facilitation of a shift from habitual to compulsive alcohol use [69, 70]. The drinking-in-the-dark and two-bottle choice drinking paradigms leveraged in the current study may not have provided a long enough alcohol exposure or induced high enough blood ethanol concentrations (BECs) to substantially recruit DLS involvement over these behaviors. DID decreases GABAergic transmission in the DLS, as measured by a decrease in mIPSC frequency, but does not alter DLS SPN spine density or spontaneous glutamatergic transmission [71]. In contrast, dendritic hypertrophy has been observed in DLS neurons along with CB1 receptor-mediated long-term depression following chronic intermittent alcohol vapor exposure, a paradigm which induces nearly twice the BEC level of DID, highlighting that some DLS alcohol drinking-induced neuroadaptations are only observed following much higher levels of alcohol exposure [72]. Therefore, it is possible that DLS astrocytes may modulate alcohol-drinking under higher alcohol-exposure conditions. Additionally, recent work has shown that the use of CalEX to inhibit astrocyte calcium signaling in the cortex decreases 2BC drinking [11], and mice overexpressing CalEX in nucleus accumbens show increased aversion-resistant drinking following a two-bottle choice drinking history [47]. Together, these results highlight the circuit-specific roles of astrocytes in alcohol stimulation and voluntary alcohol consumption, which is congruent with prevailing theories of the development of alcohol use disorder [73].

Astrocyte calcium activity is thought to in part signal behavioral state information and accordingly provides contextual modulation of circuit-specific neuronal activity and behavioral output [10, 74]. Locomotion in the open field is partly driven by a drive to explore the environment. Interestingly, we found astrocyte CalEX expression to have minimal effects on baseline locomotion (Figure 2E), yet alcohol delivery revealed a hyperlocomotion phenotype (Figure 2F, G). This is unlike that observed in the medial entorhinal cortex where astrocyte CalEX expression reduced baseline exploratory locomotion [75]. These findings suggest that astrocyte contribution to exploratory behavior is both circuit and context specific. Interestingly, our work also suggests a strong dependence of astrocyte behavioral contributions on the neuromodulatory landscape even within the same circuit. In previous work, we found that augmenting DLS astrocyte calcium signaling with chemogenetic stimulation in a mouse model of Parkinson’s disease increased open field locomotion, but only during the early phase when exploratory drive is high [40]. Hence, coordinating exploratory behavior under different contexts may be a key function of DLS astrocytes. At the same time, it is perplexing that opposing manipulations of the same cell type in the same circuit produces the same results – increased locomotion. There are pronounced differences in the striatal neuromodulatory landscape under alcohol and parkinsonian conditions. While alcohol is expected to increase dopamine release [76], in parkinsonian animals dopamine release is severely curtailed. In contrast, alcohol decreases ACh release (Figure 4), whereas dopamine loss is associated with increased DLS cholinergic signaling [77]. Collectively, these results suggest that the impact of astrocyte calcium signaling on behavioral output is not static but rather highly context-dependent and shifts with neuromodulatory conditions in different contexts. A noted limitation of the current research is that we only assessed astrocyte calcium signaling, but it is known that astrocytes use diverse additional second messengers for communication with neurons [78]. Future work will assess how additional astrocytic signals, such as cyclic adenosine monophosphate (cAMP) and adenosine, are impacted by alcohol.

Astrocytes express a variety of neurotransmitter receptors, which allows them to “listen in” to the neuronal activity around them and respond with elevations in intracellular astrocyte calcium [54]. Because astrocyte activity reflects an integration of local neuronal and neuromodulatory signals, it was important to record local DLS neuronal activity to get a more complete view of alcohol effects in the DLS circuit. We found that indirect pathway SPN activity was not significantly impacted by acute alcohol at any dose tested (Figure 3F-J), but we observed a reduction in the prominence of calcium events recorded from direct pathway neurons (Figure 3E). This reduction could reflect the astrocyte calcium inhibition that we observed. Alternatively, it is possible that both cell types show inhibition because of alcohol’s effect on an input common to both cell types. Following this rationale, we recorded neuromodulator release activity from the DLS during alcohol exposure. Previous work has shown that alcohol stimulation is under dorsal striatal control and heavily influenced by dopaminergic signaling [25]. One goal of the current work was to assess if other striatal neuromodulators contribute to the alcohol-induced stimulant response. Cerebellar and cortical astrocytes are inhibited by alcohol via inhibition of norepinephrine release [34]. To assess a contribution of NE to astrocyte calcium inhibition in the DLS, we recorded NE release from the DLS and BNST (a control brain region chosen because it has strong norepinephrine inputs [49]). During tail suspension recordings, the BNST, but not DLS, showed robust NE release (Figure S6F - H1). These findings confirmed that the DLS has a low NE-tone and suggested a striatum-specific mechanism for inhibition of astrocyte calcium activity by alcohol.

We found that acute alcohol inhibited DLS ACh release (Figure 4D - G). This finding is similar to other recent results demonstrating that alcohol decreases the spontaneous firing rate of DMS CINs in ex vivo brain slices and reduces ACh release in vivo as measured using fiber photometry with the iAChSnFR sensor [63]. Astrocytes express both nicotinic [79] and muscarinic ACh receptors (AChRs) [80, 81], which support astrocyte calcium activity [82, 83]. Given this relationship, we tested whether a reduction in ACh release following EtOH drives alcohol-induced astrocyte activity inhibition. We expressed the Gq-coupled excitatory chemogenetic actuator hM3D in CINs and tested how artificially increasing ACh release impacts alcohol effects on astrocyte activity. Although this manipulation augmented ACh release (Figure S7), it did not impact astrocyte calcium activity under basal or alcohol conditions (Figures 5, 6, respectively). Given that CINs are tonically active in the striatum [84], continually and locally releasing ACh, it is possible that further increasing ACh release is ineffective due to a competing saturation effect wherein we are unable to further modulate downstream impact on astrocyte activity. In future studies, it will be important to employ loss-of-function tools that decrease ACh release to test this possibility. Notwithstanding these concerns, our current experiments rule out one direction of the potential interaction between ACh release and astrocyte calcium. We have not addressed the possibility that alcohol’s inhibition of astrocyte calcium signaling drives the inhibition of ACh release. Recent work shows that optogenetic activation of striatal astrocytes results in the modulation of CIN activity [62], suggesting that this option is not only possible but may even be likely. It is also possible that the observed alcohol-induced suppression of ACh release and astrocyte calcium inhibition are unrelated and that astrocytes and CINs independently regulate alcohol-related behaviors. More work is needed to establish the exact relationship between alcohol stimulation, striatal astrocyte calcium activity and ACh release.

We found an interesting trend where mice showed lower alcohol stimulation with chemogenetic CIN stimulation (Figure S8), suggesting that striatal ACh release curtails alcohol stimulation. These findings also suggest that the EtOH-induced suppression of ACh release that we observed contributes to alcohol stimulation. Our collective findings are broadly consistent with the idea that CIN activity during alcohol exposure controls the level of alcohol-induced stimulation and astrocytes “fine tune” the response. CalEX expression did not lead to overall higher alcohol-induced stimulation but potentiated the stimulant response at the moderate dose of 1g/kg (Figure 2), suggesting a modulatory role. Moreover, CalEX expressing astrocytes are presumably already inhibited, which would preclude further inhibition by alcohol. Since enhanced alcohol stimulation observed under these conditions is unlikely to be driven by further inhibition of astrocyte calcium, these findings suggest that the residual astrocyte activity observed after alcohol delivery in our fiber photometry experiments plays a protective role against excessive stimulation. In this scenario, when astrocyte calcium signaling is inhibited by CalEX, these cells are unavailable to adaptively fine tune the neuronal processes which govern alcohol stimulation and as a result mice become more susceptible to producing a maladaptive behavioral response (i.e., a greater alcohol-induced stimulation). It is well established that acute alcohol stimulates striatal dopamine release and that alcohol stimulation is heavily regulated by dopaminergic inputs [25, 27, 85]. Striatal dopamine-ACh dynamics are carefully co-regulated to produce complex behaviors [86, 87]. One additional hypothesis raised by the current findings is that ACh inputs serve as a gating signal, activating a permissive state which by itself doesn’t modulate calcium activity but allows astrocytes to respond to other neurotransmitters, such as dopamine, to fine tune alcohol stimulation. Recent work has shown GPCR gating of astrocyte neurotransmitter reponses, supporting this possibility [88].

In summary, we have identified that alcohol inhibits both DLS astrocyte calcium activity and ACh release. Further, we have shown that inhibition of DLS astrocytes using a molecular genetic manipulation resulted in a more pronounced alcohol-induced stimulant response, suggesting that DLS astrocytes play an important role in modulating this behavior. We have also established that SPN activity, NE release, and ACh release are unlikely to contribute to alcohol-induced astrocyte activity suppression. Our experiments uncover the multifaceted effects of acute alcohol on the neuronal, non-neuronal, and neuromodulatory elements of the DLS circuit. Given the clinical relevance of the alcohol-induced stimulant response in predicting the future development of AUD, it is critically important to understand the biological mechanisms which drive this and other risk factors for this disease.

## Funding and Acknowledgements

This work was supported by the National Institute on Alcohol Abuse and Alcoholism (NIAAA; F32-AA032173 to C.A., R01-AA030594 to R.H., and R00-AA027750 to M.B.), the National Insitute on Drug Abuse (NIDA; T32-DA055569 to C.A. and M.M.), the National Science Foundation (NSF-GRFP to N.C.B), and a Rutgers Presidential Postdoctoral Fellowship (C.A.).

We thank Angelica Vellore for assistance with histology. We also thank Drs. Veronica Alvarez and Grayson Sipe for valuable feedback on this work.

## Author Contributions

**Cherish Ardinger**: writing – original draft, conceptualization, investigation and formal analysis (fiber photometry, chemogenetic and CalEX experiments, histology and imaging), writing – review and editing, visualization; funding acquisition; **Anagha Kalelkar:** investigation and formal analysis (some fiber photometry and CalEX experiments); writing – review and editing**; Maxwell Madden:** investigation and formal analysis (ex-vivo two-photon imaging experiments), writing – review and editing; **Ananya Gunda, Arnav Patel, and George Xanthos:** investigation (histology and imaging; assistance with fiber photometry recordings, transgenic breeding colony management), writing – review and editing; **Mariam Mahboob and Ali Khawaja:** investigation and formal analysis (two-bottle choice drinking CalEX experiments); **Nancy Collie-Beard:** investigation and formal analysis (drinking-in-the-dark CalEX experiments, assistance setting up behavioral testing equipment), writing – review and editing; **Miriam Bocarsly:** supervision, funding acquisition, writing – review and editing; **Rafiq Huda:** conceptualization, supervision, funding acquisition, writing – review and editing.

## Competing Interests

The authors have nothing to disclose.

---

**Supplementary Figure 1.**
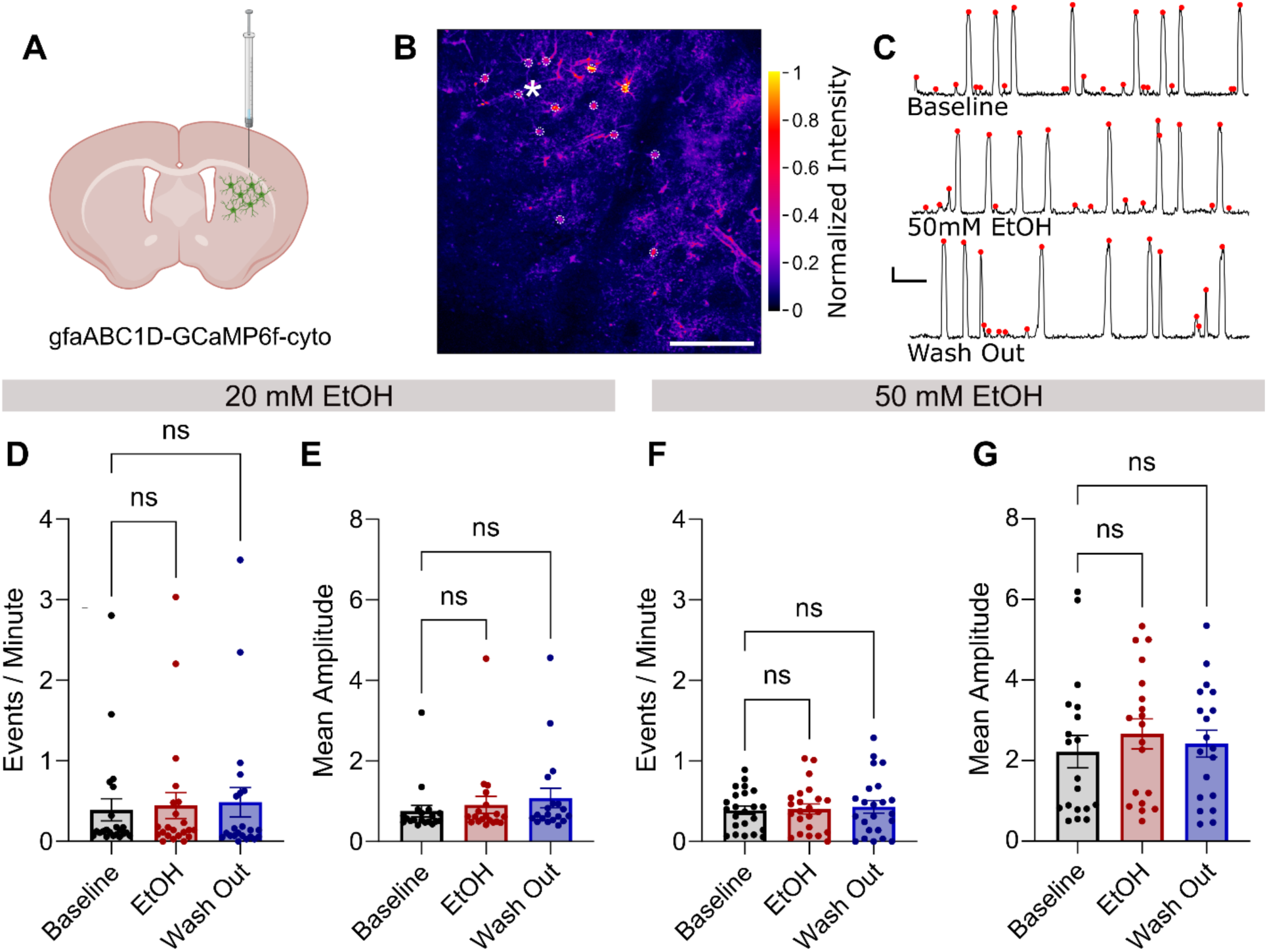
No EtOH-induced astrocyte calcium activity changes were observed in ex vivo brain slices. **A)** Mice were given a stereotaxic injection of gfaABC1D-GCaMP6f-cyto in the DLS to measure astrocytic calcium activity in acutely prepared ex-vivo brain slices. **B)** Representative 2P images (normalized maximum projection) of astrocytes expressing GCaMP6f. Astrocyte somas were manually identified and selected as ROIs for calcium imaging analysis. Scale bar is 100 μm. **C)** Representative traces of the spontaneous calcium activity of an identified ROI from B (indicated with asterisk) during the baseline, 50 uM EtOH, and wash out conditions. Scale bar is 1 min (horizontal) by 1 ΔF/F (vertical). **D)** No significant difference in events per minute was found when 20 mM EtOH was applied to the slices, *F*(1.519, 31.90) = 1.995, *p* = 0.1612, η^2^p = 0.0867 (Greenhouse-Geisser corrected repeated measures one-way ANOVA; n = 22, from 6 animals). **E)** No significant difference in mean amplitude was found when 20 mM EtOH was applied to the slices, *F*(1.209, 21.77) = 0.9882, *p* = 0.3479, η^2^p = 0.0520 (Greenhouse-Geisser correctedrepeated measures one-way ANOVA; n = 22, from 6 animals). **F)** No significant difference in events per minute was found when 50 mM EtOH was applied to the slices, *F*(1.382, 30.40) = 0.5517, *p* = 0.5183, η^2^p = 0.0244 (Greenhouse-Geisser corrected repeated measures one-way ANOVA; n = 23, from 5 animals). **G)** No significant difference in mean amplitude was found when 50 mM EtOH was applied to the slices, *F*(1.356, 24.41) = 0.7424, *p* = 0.4368, η^2^p = 0.0396 (Greenhouse-Geisser corrected repeated measures one-way ANOVA; n = 23, from 5 animals). All error bars represent standard error of the mean (SEM).

**Supplementary Figure 2.**
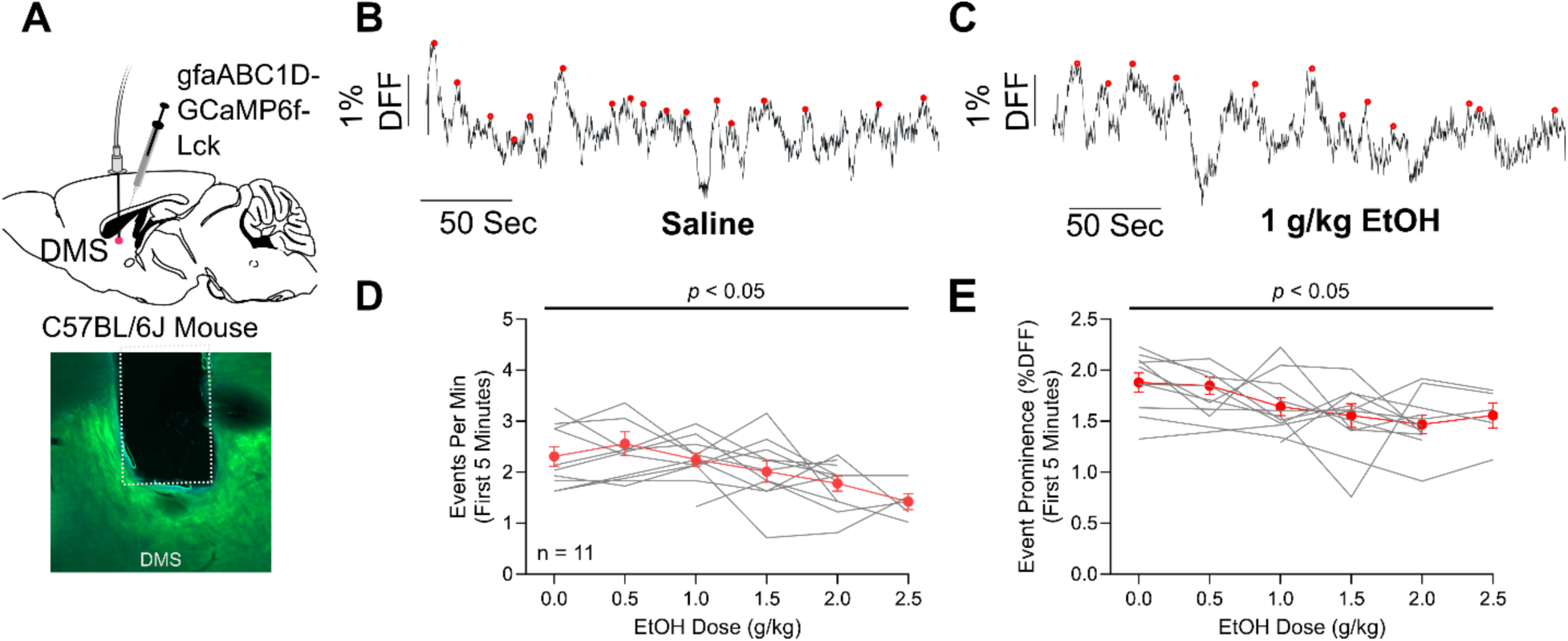
Acute alcohol inhibits DMS astrocyte calcium signaling in vivo. **A)** To record DMS astrocyte calcium activity, mice (n = 11) were given a stereotaxic injection of a GCaMP virus with an astrocyte-specific promoter with accompanying fiber optic implant for in-vivo fiber photometry calcium recording (top). Representative viral expression is shown (bottom). **B)** Example astrocyte calcium traces when mice are given saline or **C)** 1 g/kg EtOH. **D)** Acute alcohol inhibits astrocyte calcium event frequency; *F*(5, 36) = 2.989, *p* = 0.023 (repeated measures mixed effects analysis), and **E)** decreases event prominence; *F*(5, 35) = 2.851, *p* = 0.0291 (repeated measures mixed effects analysis). All error bars represent standard error of the mean (SEM).

**Supplementary Figure 3.**
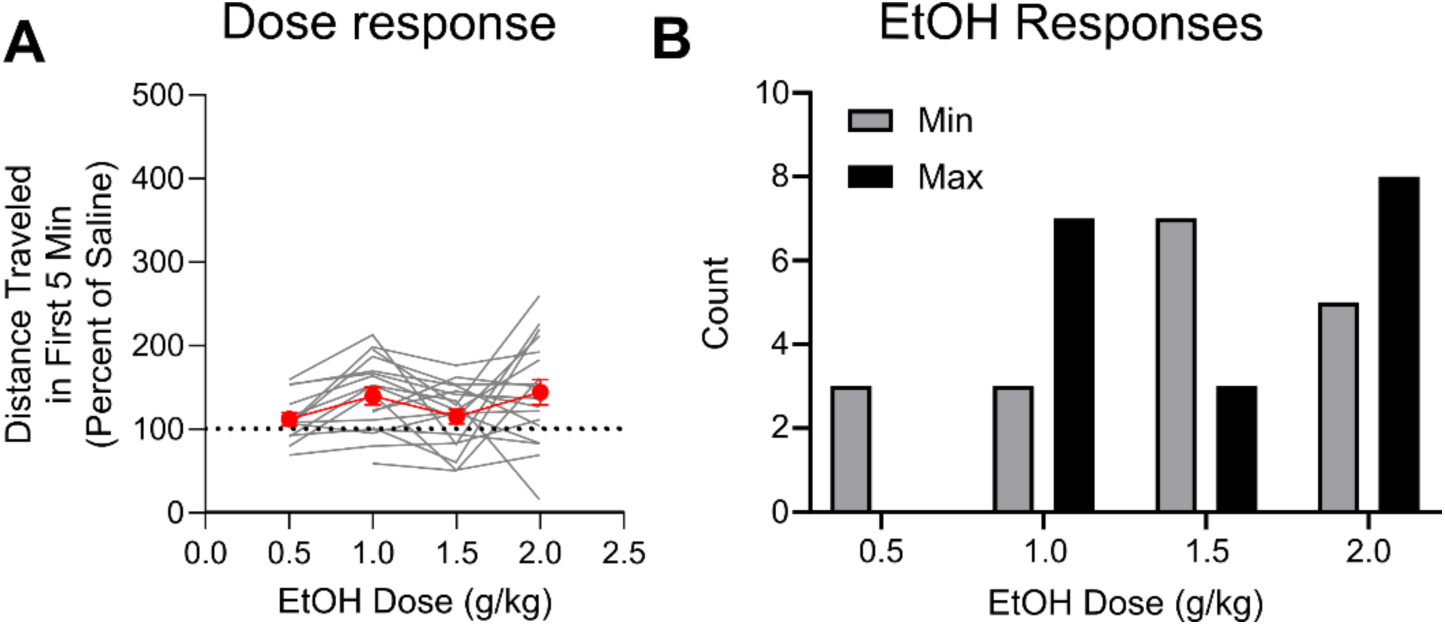
Mice display variability in their stimulant response to alcohol during in vivo fiber photometry. **A)** Distance travelled across EtOH doses in the first 5 minutes expressed as a percentage of saline control from DLS (n = 7) and DMS (n = 11) striatal astrocytic calcium recordings. No significant differences were noted across doses, *F*(1.895, 29.05) = 2.811, *p* > 0.05 (repeated measures mixed effects analysis), but all doses produced a mean response greater than 100% of saline, suggesting some level of EtOH-induced stimulation. **B)** Across mice there was variability in the EtOH dose that produced the minimal and maximal locomotion response (normalized to saline); Fisher’s exact test comparing minimum and maximum doses: *p* = 0.0834.

**Supplementary Figure 4.**
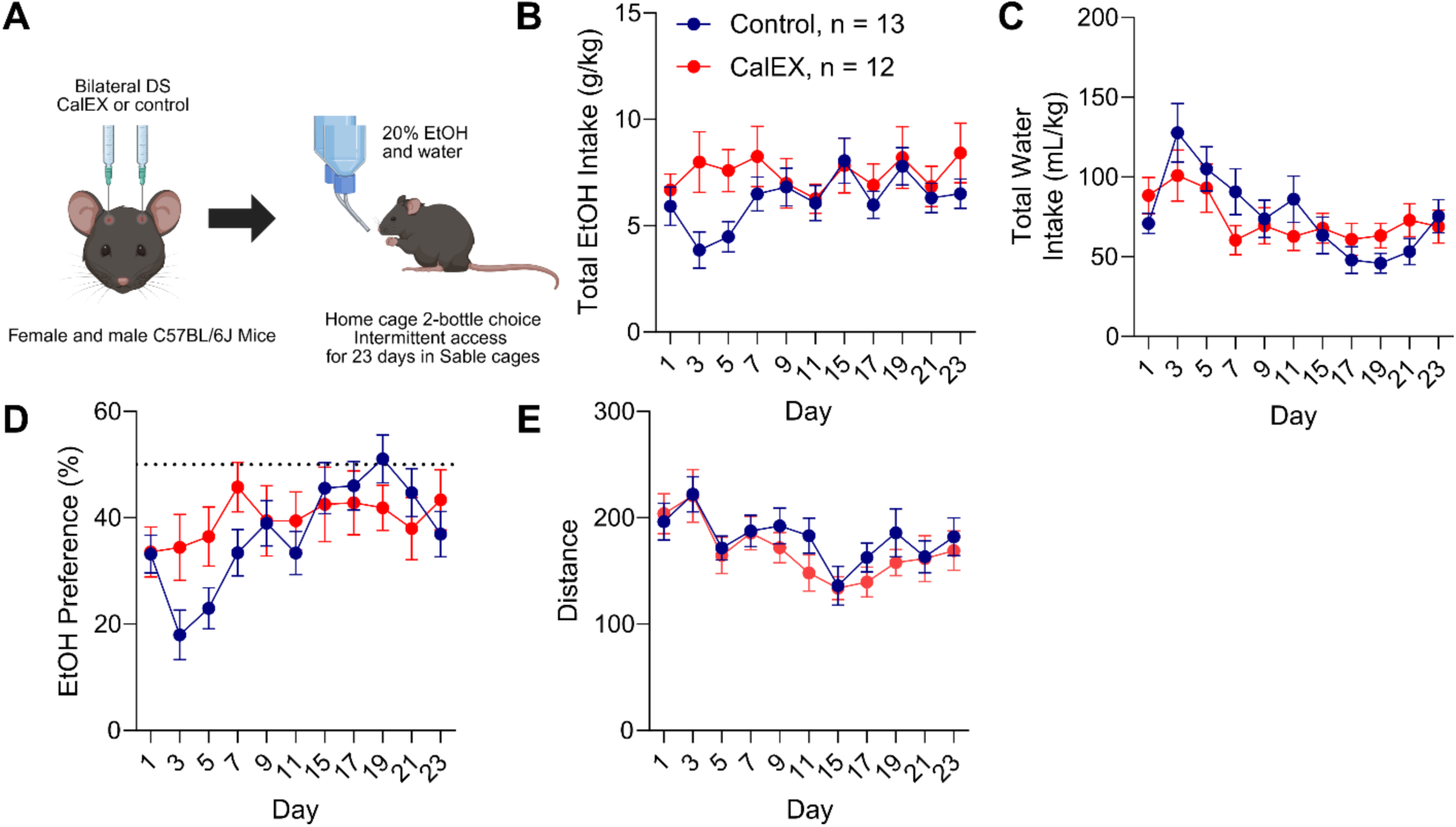
Effect of astrocyte calcium activity inhibition on home cage 2-bottle choice drinking. **A)** C57BL/6J mice received bilateral viral injections to express CalEX or a control fluorophore in DLS astrocytes. After recovery, mice performed 23 days of intermittent access 2-bottle choice (2BC) drinking (20% EtOH) in Sable cages. **B)** Across 2BC testing days, there were no differences in total daily EtOH intake, main and interaction effects of day and virus all *p* > 0.05 (repeated measures mixed-effects analysis). **C)** Analysis of water intake indicated a significant main effect of day, *F*(4.595, 104.8) = 8.916, *p* < 0.0001, and a trend for a day x virus interaction, *F*(4.595, 104.8) = 2.280, *p* = 0.057 (Greenhouse-Geisser corrected repeated measures mixed-effects analysis). **D)** Analysis of EtOH preference indicated a main effect of day, *F*(6.269, 142.3) = 4.814, *p* = 0.0001, where EtOH preference in both viral groups generally increased over days (Greenhouse-Geisser corrected repeated measures mixed-effects analysis). **E)** Analysis of distance traveled indicated a main effect of day, *F*(5.170, 118.9) = 10.81, *p* < 0.0001, η^2^p = 0.319, where distance in both viral groups steadily decreased over days (Greenhouse-Geisser corrected repeated measures 2-way ANOVA). All error bars represent standard error of the mean (SEM).

**Supplementary Figure 5.**
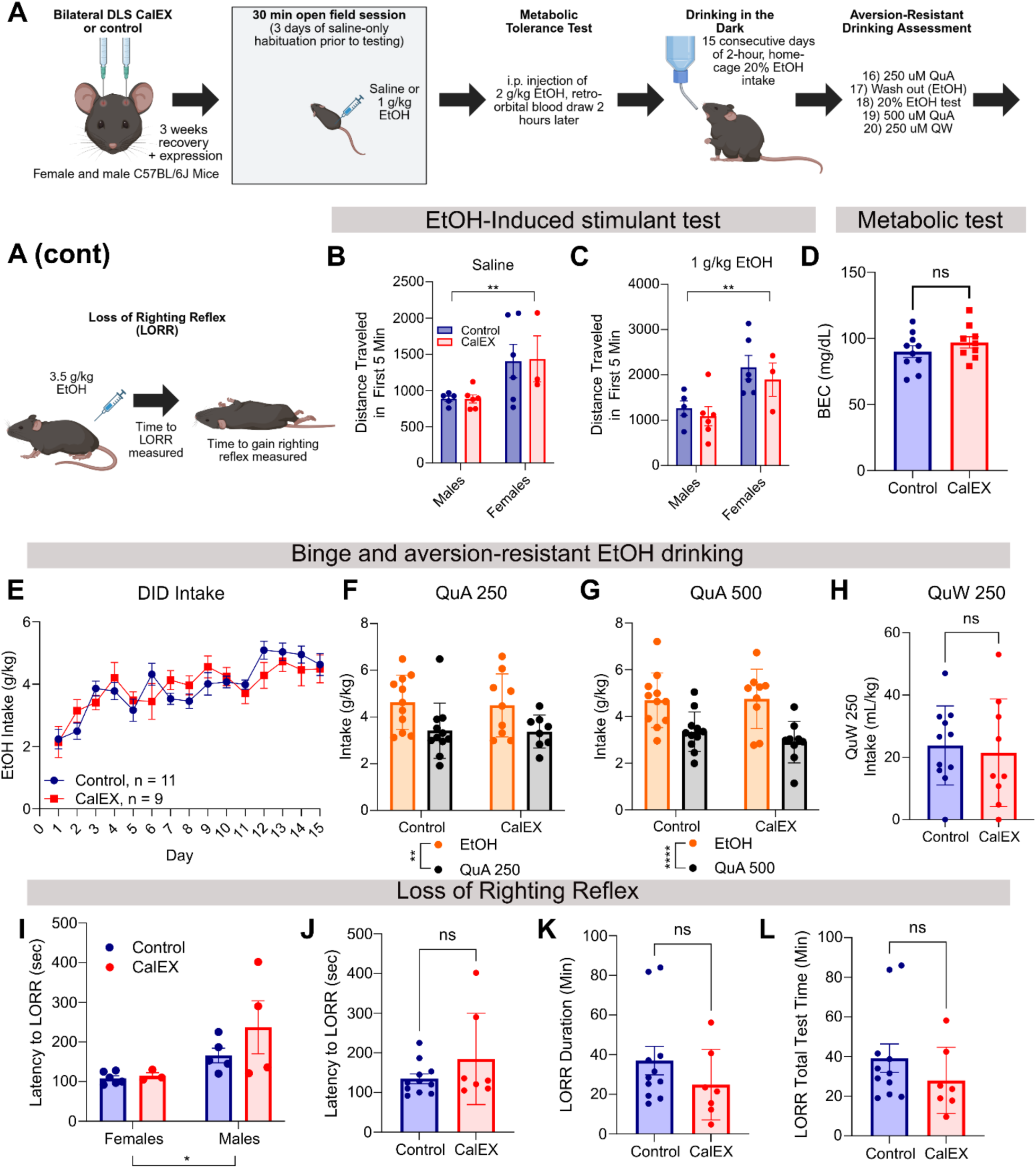
Effect of DLS astrocyte calcium activity inhibition on binge alcohol drinking, home-cage aversion-resistant alcohol drinking, and loss of righting reflex. **A)** C57BL/6J mice bilaterally expressing CalEX or a control fluorophore in DLS astrocytes underwent a sequence of EtOH-related behavioral testing. **B)** Female mice moved more in the first 5 min of open field testing than males when given saline regardless of viral condition, significant main effect of sex; *F*(1, 16) = 9.083, *p* = 0.0082, η^2^p = 0.362 (two-way ANOVA). **C)** The same was found when mice were given 1 g/kg EtOH, main effect of sex; *F*(1, 16) = 11.45, *p* = 0.0038, η^2^p = 0.272 (two-way ANOVA). **D)** BECs did not differ between viral groups, indicating that CalEX and control mice metabolize EtOH at a similar rate, *t*(17) = 1.135, *p* = 0.272 (unpaired t-test). **E)** Viral groups drank similar amounts of EtOH during DID testing: no significant main effect of virus, *F*(1, 18) = 0.004891, *p* = 0.945; or interaction of day x virus, *p* > 0.05. A main effect of day indicated that both viral groups generally increased DID EtOH intake over days, *F*(7.387, 133.0) = 11.11, *p* <0.0001, η^2^p = 0.381 (Greenhouse-Geisser corrected two-way ANOVA). **F)** When mice were presented with quinine-adulterated alcohol (250 µM; QuA 250), both viral groups significantly decreased their intakes as compared to the last day of non-adulterated DID, indicating that neither group is displaying aversion-resistant EtOH intake; main effect of solution: *F*(1, 17) = 14.83, *p* = 0.0013 (two-way repeated measures mixed effects analysis; note one male CalEX mouse had a leaky sipper on the QuA test day and was excluded on this day). **G)** The same pattern was found with a higher concentration of quinine-adulterated EtOH (QuA 500); main effect of solution: *F*(1, 18) = 26.93, *p* <0.0001, η^2^p = 0.599 (two-way ANOVA). **H)** Viral groups are equally able to taste the 250 µM quinine when it is in water only, *t*(14.40) =0.3387, *p* = 0.7397. **I)** Male mice took longer to lose righting reflex, significant main effect of sex: F (1, 14) = 7.688, *p* = 0.0150, η^2^p = 0.354. Of note, two calEX male mice never lost righting reflex following the 3.5 g/kg EtOH injection (were monitored for > 1 hour, which is why there are only 4 M CalEX mice represented on the graph). This phenomenon was not observed in any other group. **J)** When collapsed on sex (as there was no sex x virus interaction), viral groups did not differ in latency to LORR, *t*(16) = 1.336, *p* = 0.2003 (unpaired t-test). **K)** Similarly, there was no difference in LORR duration, *t*(16) = 1.142, *p* = 0.2701, or **L)** total LORR test time, *t*(16) = 1.084, *p* = 0.2943. All error bars represent standard error of the mean (SEM).

**Supplementary Figure 6.**
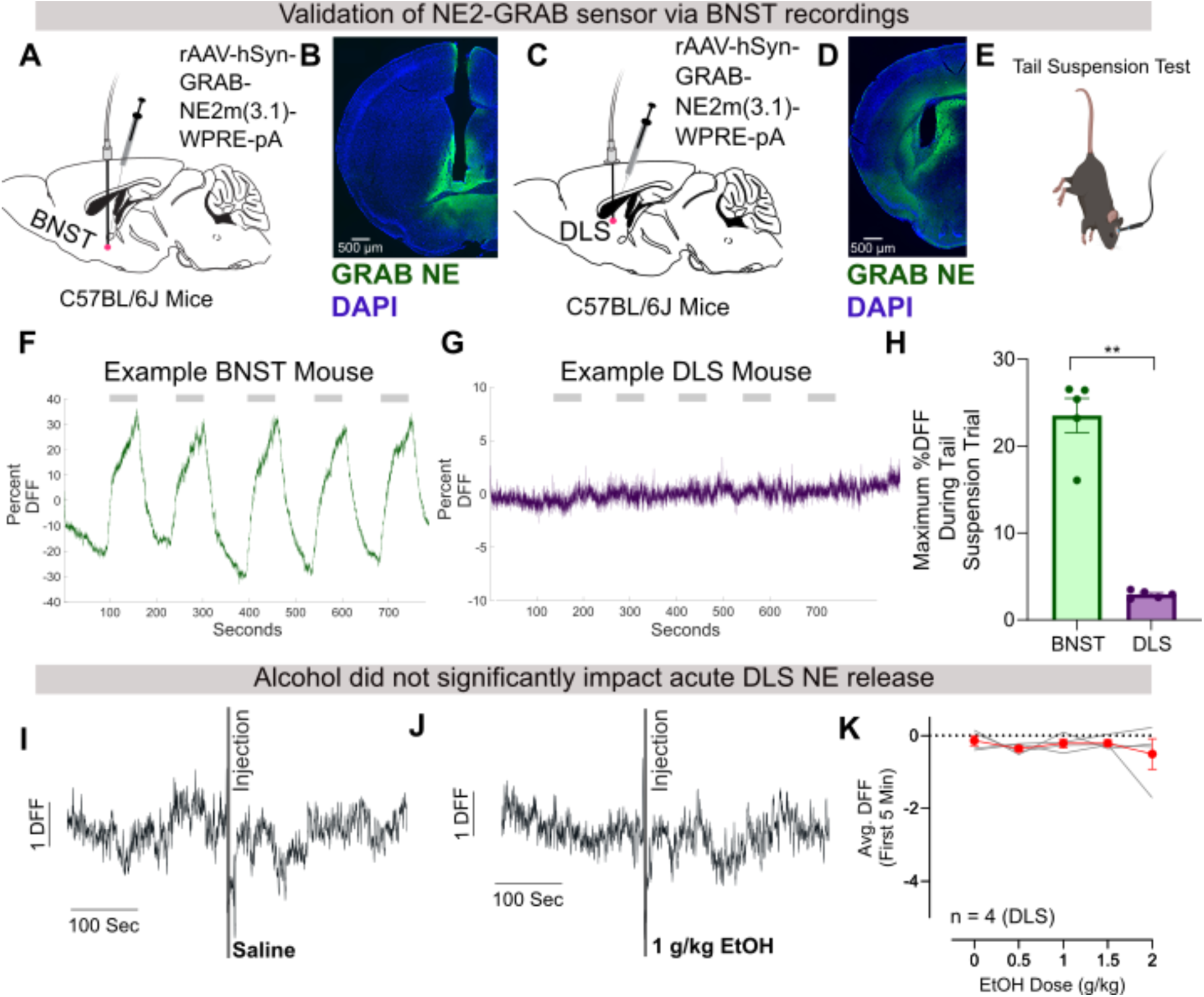
GRAB-NE sensor validation indicates that the DLS has a low NE-tone; acute alcohol does not impact NE release in the DLS. **A)** C57BL/6J mice received a stereotaxic injection of GRAB-NE 3.1 with optical fiber implant in the BNST. **B)** Representative image of BNST GRAB-NE viral spread and fiber placement. **C)** Separate mice received GRAB-NE and a fiber in the DLS. **D)** Representative image of DLS GRAB-NE viral spread and fiber placement. **E)** Mice underwent 5 tail suspension trials during fiber photometry recordings to elicit NE release. Mice were suspended for one minute with at least one minute of rest before the next suspension trial. **F)** An example trace of NE release in the BNST during tail suspension trials (gray shadings). **G)** Example trace of NE release in the DLS during tail suspension trials. **H)** Recordings from the BNST indicate significantly higher max percent DFF during tail suspension trials than from the DLS, *F*(1, 5) = 20.34, *p* = 0.0063, η^2^p = 0.778 (two-way ANOVA; brain region x trial). **I)** Example DLS NE DFF trace when a mouse received saline, or **J)** 1 g/kg EtOH. **K)** Across EtOH doses, there was no significant change in first 5-minute avg. DFF, *F*(1.385, 4.154) = 0.4871, *p* = 0.5819 (Greenhouse-Geisser corrected repeated-measures one-way ANOVA). All error bars represent standard error of the mean (SEM).

**Supplementary Figure 7.**
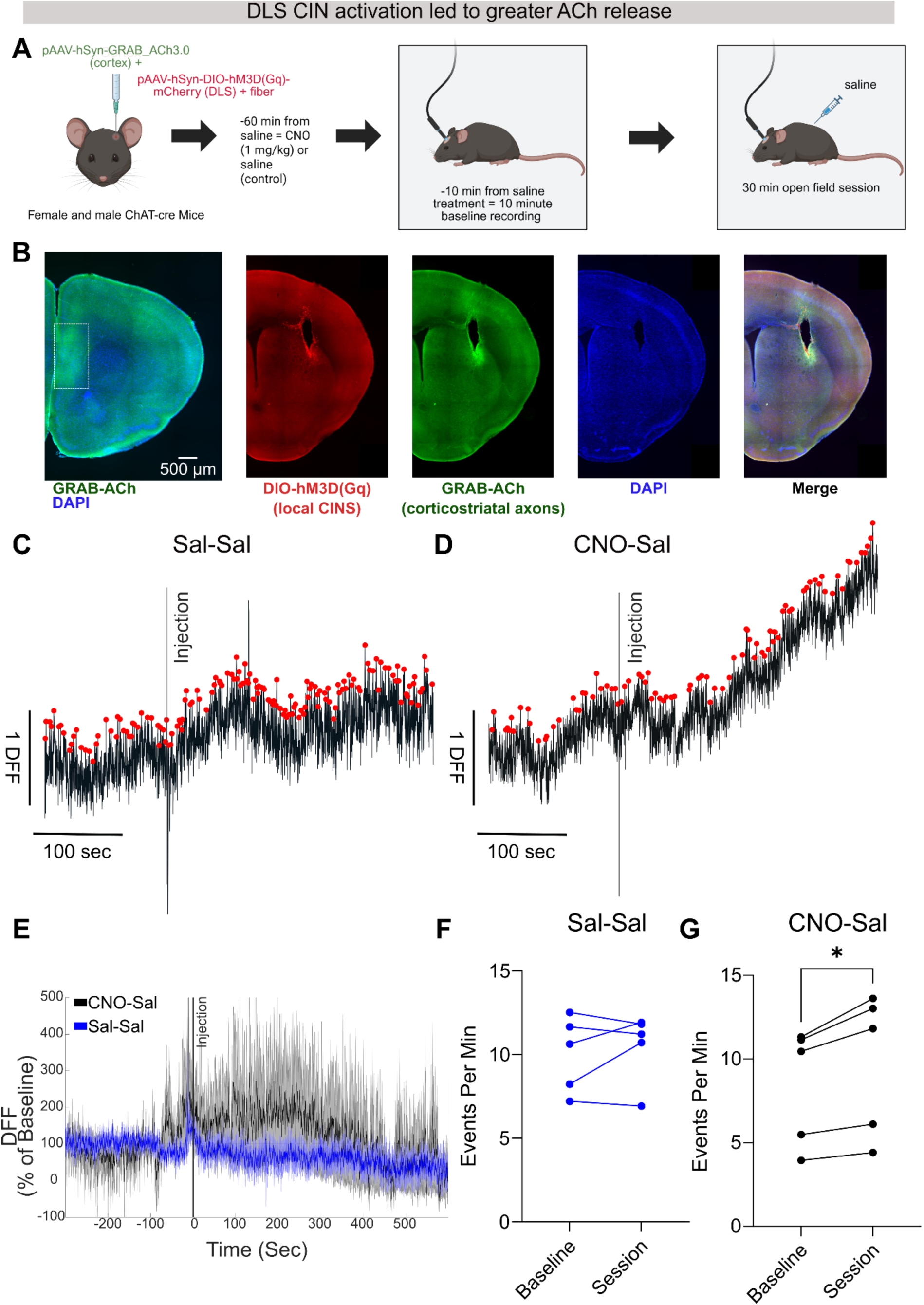
Chemogenetic stimulation of DLS CINs resulted in greater ACh release as measured by the ACh-GRAB sensor. **A)** ChAT-Cre mice received a stereotaxic injection of GRAB-ACh3.0 in the cortex. In a second surgery, mice received a stereotaxic injection of a Cre-dependent excitatory DREADD and an optical fiber implant in the DLS. Following recovery, mice received CNO or saline pre-treatment (order of treatment was counterbalanced across subjects). A 10-minute baseline recording was acquired 50 minutes after the pretreatment. Then a saline injection was given, followed by a 30-minute open-field fiber photometry recording to assess extracellular ACh release. **B)** Representative image of GRAB-ACh (green) and DAPI (blue) expression in the cortex (far left). The white dotted box indicates the viral GRAB-ACh expression; representative images of DREADD expression (red), GRAB-ACh expression in corticostriatal axonal projections, DAPI, and merged composite image in the DLS. **C)** Representative GRAB-ACh traces for saline-saline and **D)** CNO-saline conditions. The vertical line labeled injection indicates the point at which the ‘baseline’ recorded ended and the mice received the saline treatment and subsequent 30-minute open-field session recording. **E)** Mean DFF traces for Sal-Sal and CNO-Sal condition (expressed as a percent of baseline). **F)** There was no significant difference in the number of detected ACh events when comparing the pre-injection baseline and post-injection session when mice were given saline pre-treatment, *t*(4) = 0.7680, *p* = 0.4853 (paired t-test); note that the first 10 minutes of the full 30 minute session was used for post-injection data to ensure that the same amount of time is considered for both recordings. The same applies to G. **G)** Mice displayed significantly more ACh events following the saline injection when given a CNO pre-treatment, *t*(4) = 3.750, *p* = 0.0200 (paired t-test).

**Supplementary Figure 8.**
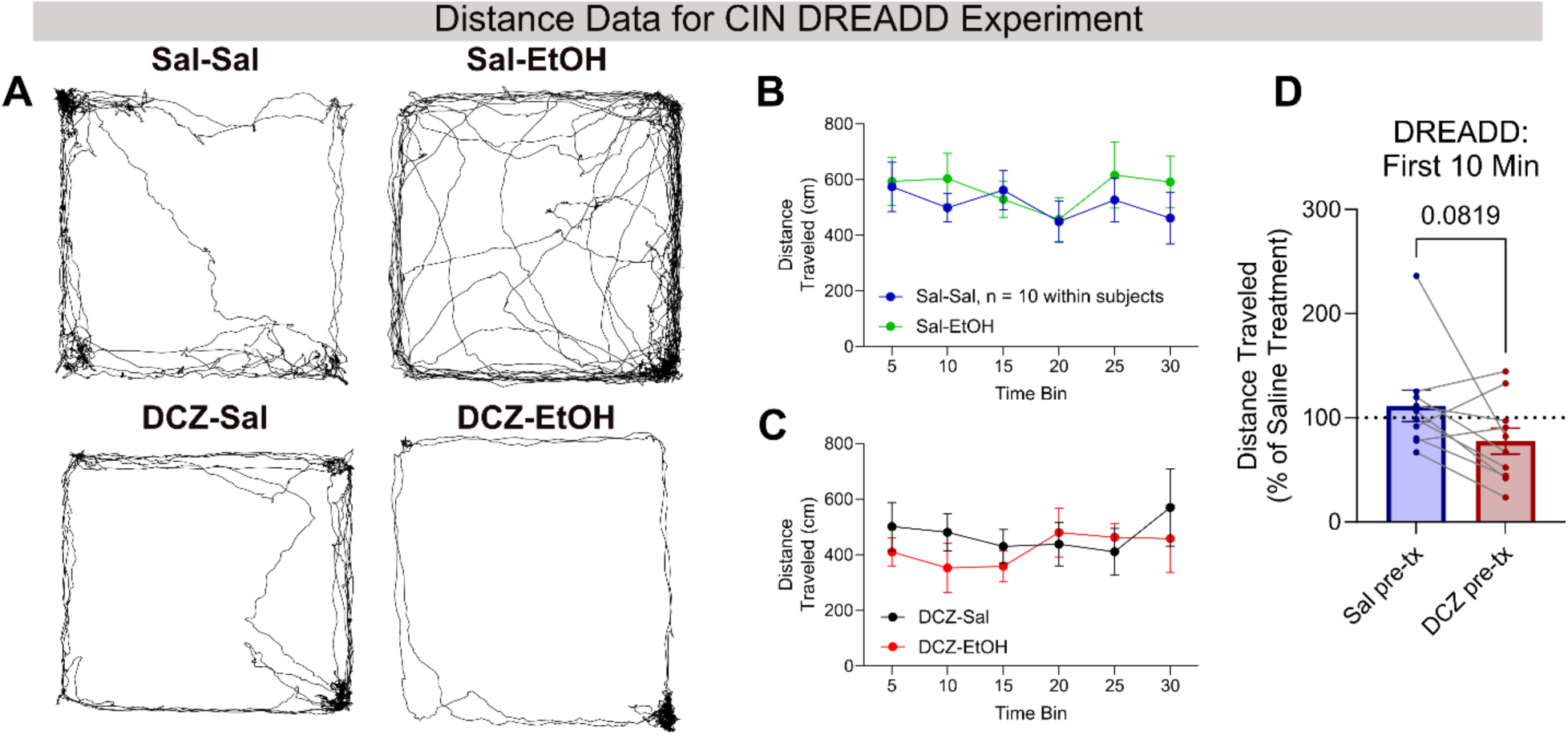
Activating cholinergic interneurons shows a trend for blunting the alcohol-induced stimulant response. **A)** 2D track plots showing examples of locomotion with the four treatment conditions. **B)** Across the 30-minute open field session, there was no significant difference in movement between sal-sal and sal-EtOH treatment, *F*(1, 18) = 0.3723, *p* = 0.5494 (repeated measures two-way ANOVA; treatment x time bin). **C)** Similarly, there was no difference between DCZ-sal and DCZ-EtOH treatment, *F*(1, 18) = 0.5309, *p* = 0.4756 (repeated measures two-way ANOVA; treatment x time bin). **D)** Distance traveled in the first 10 minutes was normalized to within-treatment saline (i.e., saline-EtOH was normalized to saline-saline and DCZ-EtOH was normalized to DCZ-saline) to facilitate comparison across pre-treatment conditions. This analysis identified a trend where DCZ pre-treatment blunted the alcohol-induced stimulant response, *t*(9) = 1.958, *p* = 0.0819, *d* = 0.7735 (paired t-test). All error bars represent standard error of the mean (SEM).

**Supplementary Table 1.**
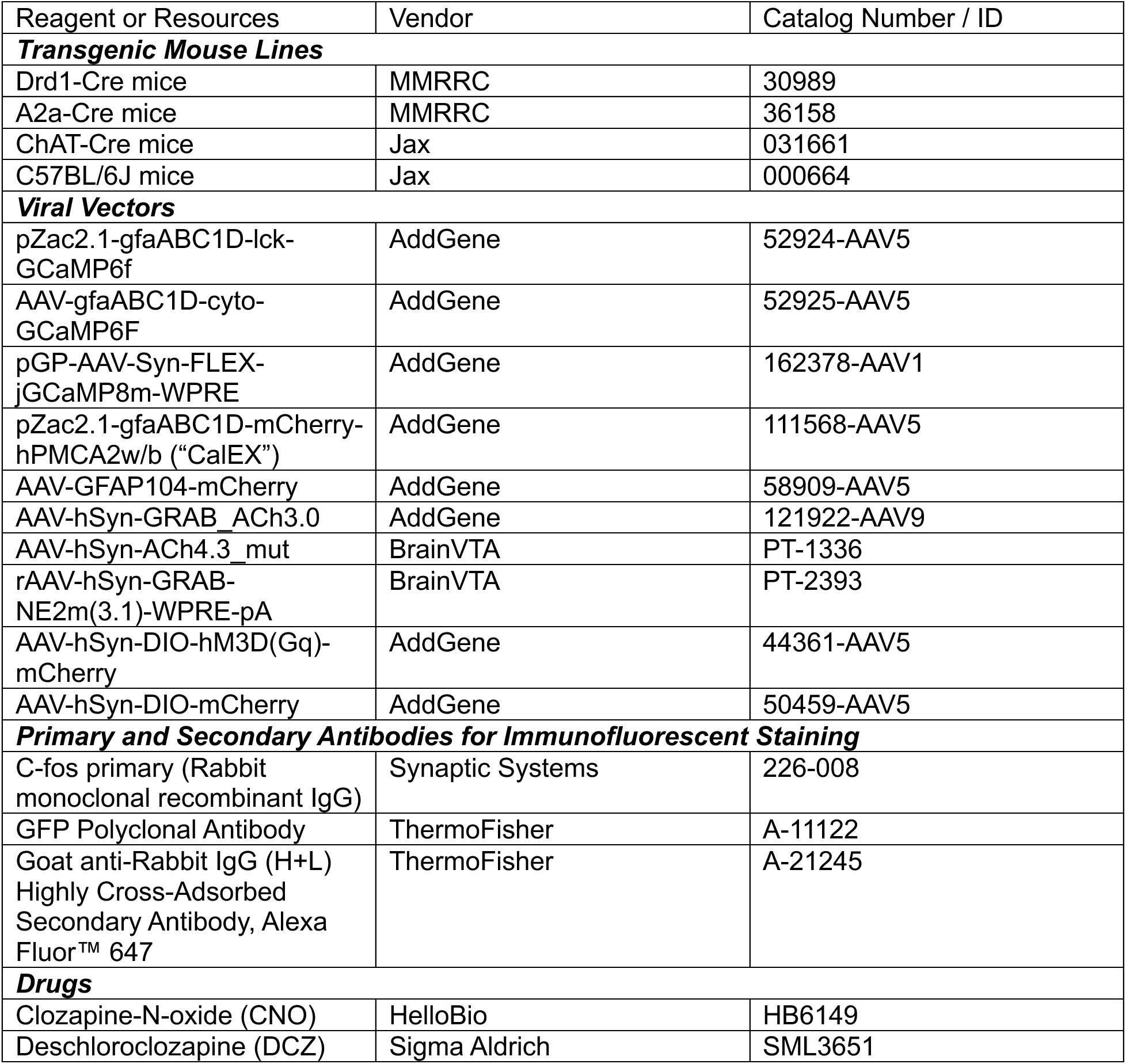
Key Reagents and Resources.

**Supplementary Table 2.**
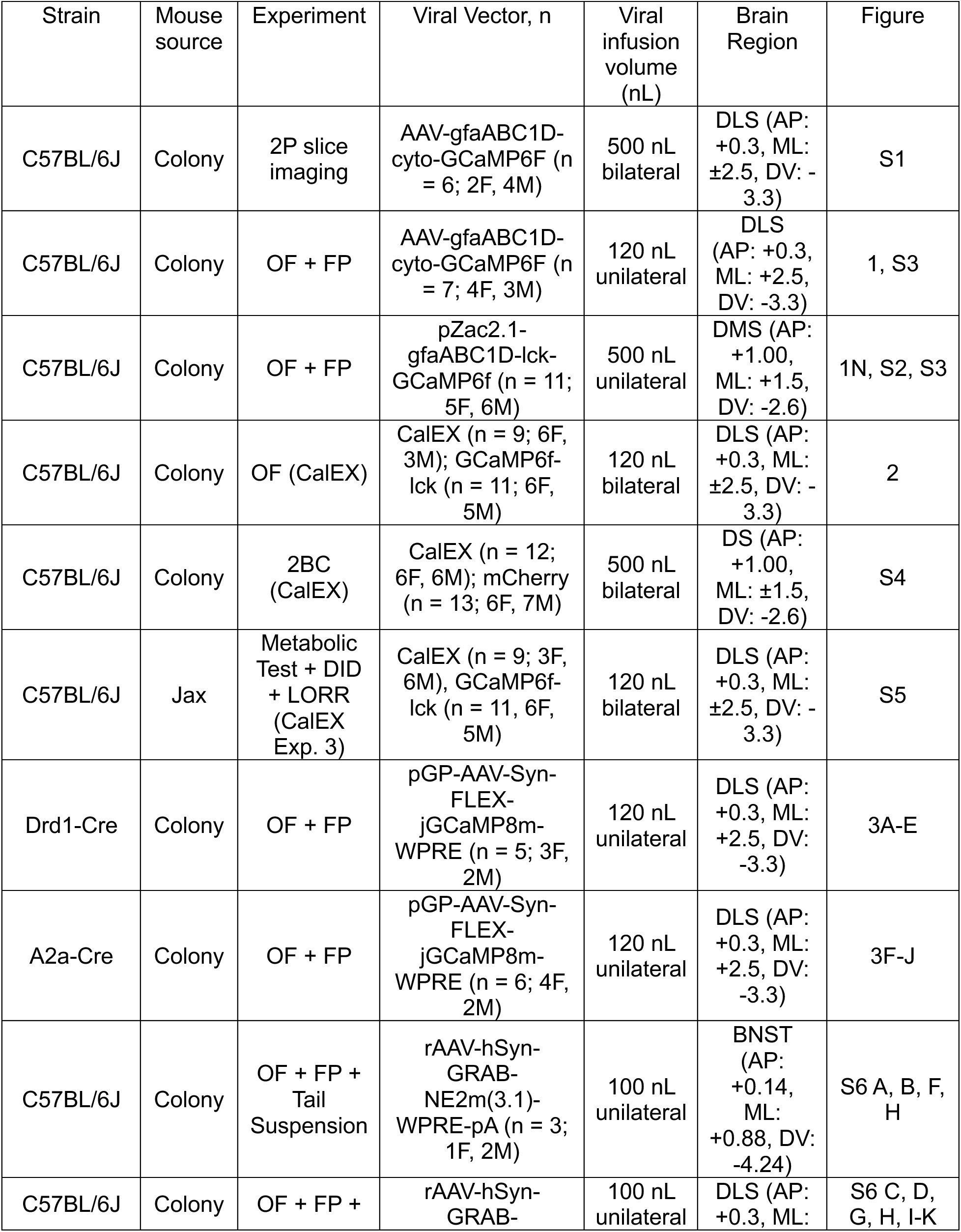
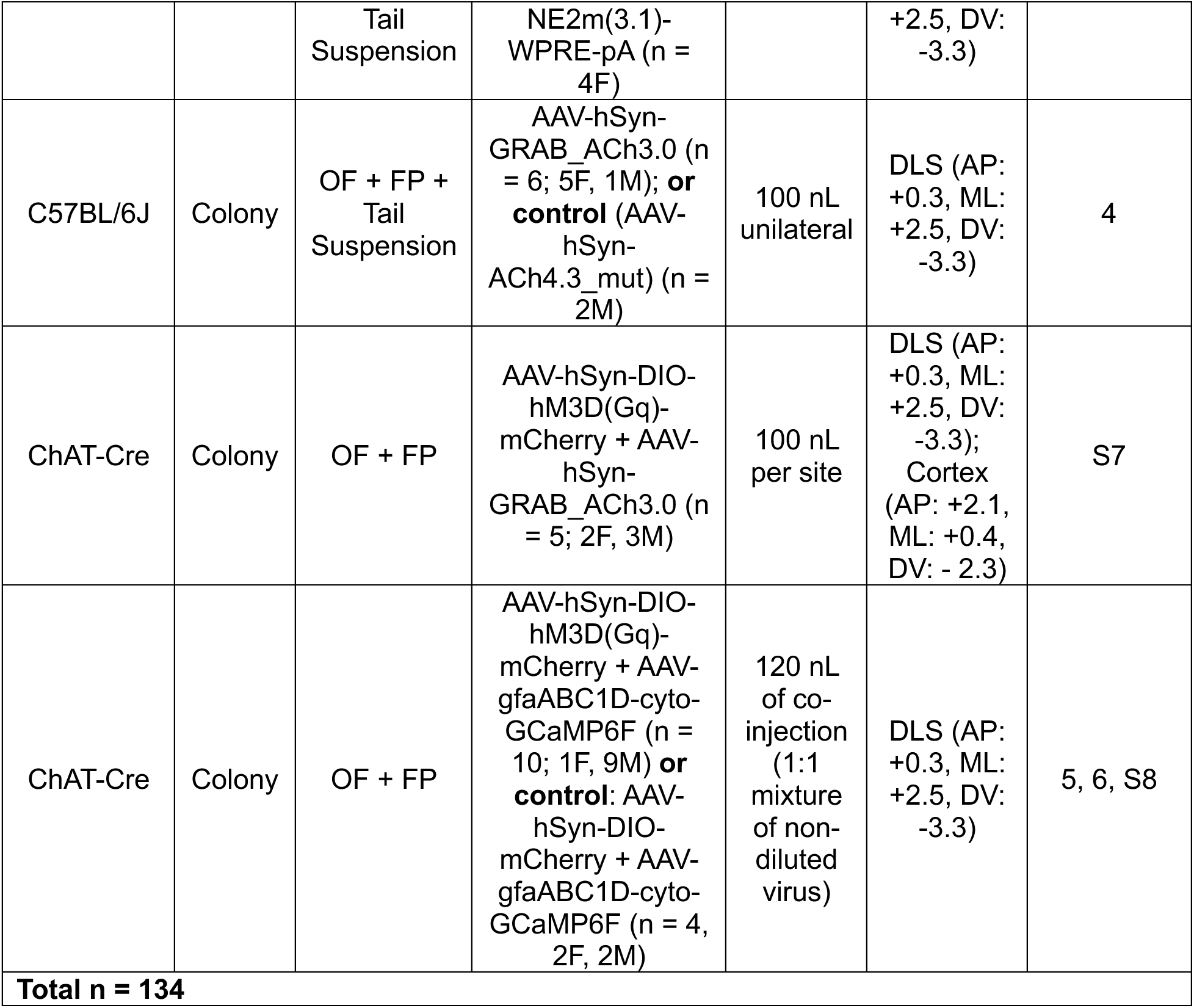
Overview of Subjects.

**Supplementary Table 3.**
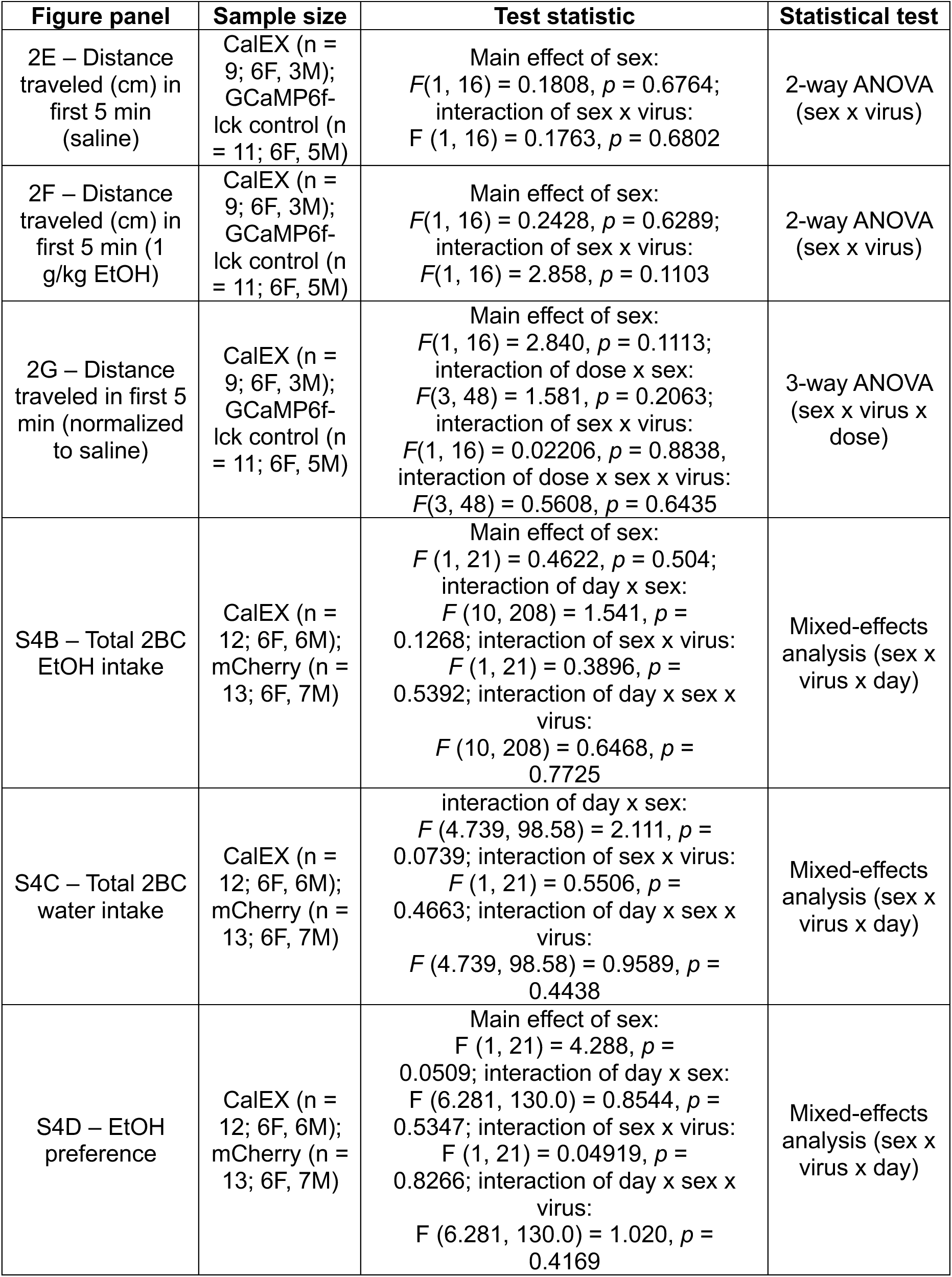
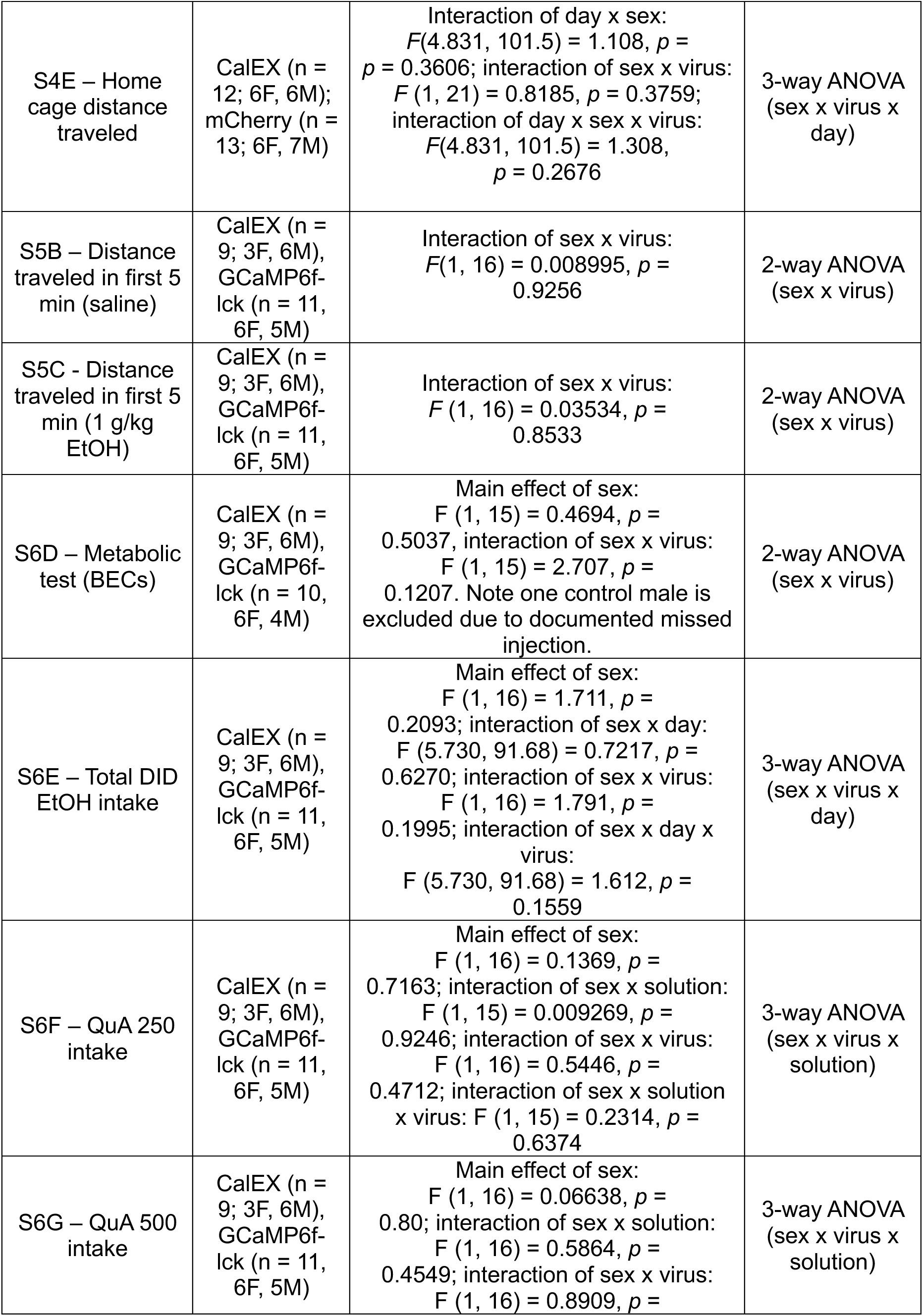
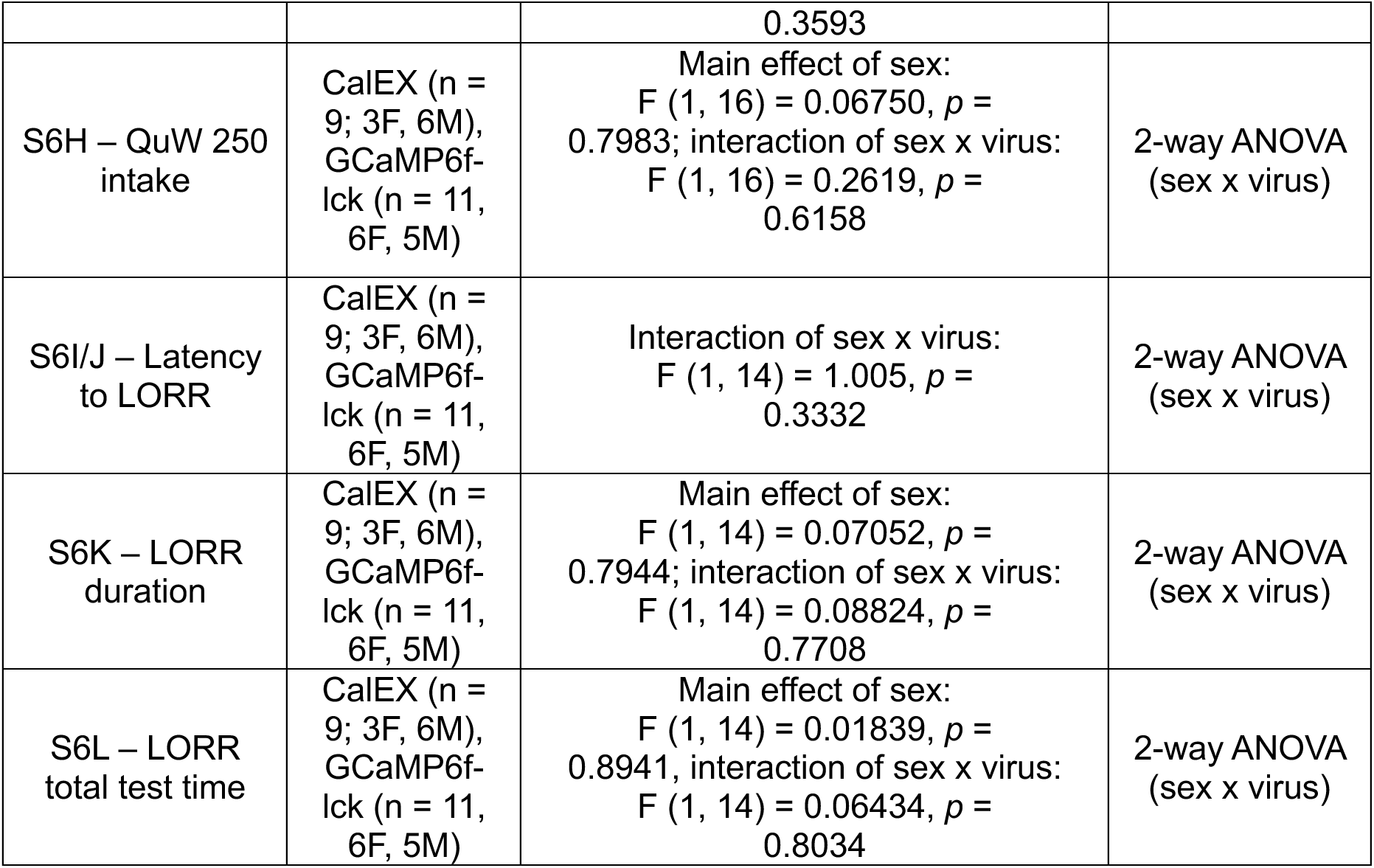
Null sex effects in behavioral experiments.

## Supplementary Note 1

### Extended Methods

#### CalEX: 2-bottle choice (2BC) home-cage alcohol drinking

Previous work has shown that the stimulant response to alcohol predicts the future development of alcohol use disorder [18, 20, 21]. Therefore, we were also interested in assessing if striatal astrocytes modulated other alcohol-related behaviors, such as home-cage voluntary alcohol drinking. For our second CalEX experiment, we assessed if there was a role for striatal astrocytes in modulating voluntary alcohol consumption. A new group of C57BL/6J mice bilaterally expressing CalEX (n = 12; 6F, 6M) or a control viral fluorophore (n = 13; 6F, 7M) in the dorsal striatum (AP: +1.00, ML: +1.5, DV: -2.6) received home-cage access to water and 20% EtOH (diluted from 190 proof EtOH) administered via Promethion Rodent Behavioral Phenotyping Cages (Sable Systems International). Mice were single housed and received intermittent access to EtOH and water for 24 hours every other day for a total of 23 days. On non-testing days, mice only had access to water. Mice had ad libitum access to chow during testing. Using these phenotyping cages provided the ability to record intake of either solution with high temporal and spatial resolution. Following testing in the Sable Systems cages, mice subsequently underwent 2BC “traditional” intermittent access 2BC testing for 10 days in the home cage with standard drinking tubes created from 50mL tubes outfitted with sippers and rubber stoppers following published protocols [89].

#### CalEX: EtOH metabolism test

For our final CalEX experiment, mice underwent a sequence of testing EtOH-related behaviors, which is outlined in Figure S5A. In this last CalEX experiment, C57BL/6J mice bilaterally expressing CalEX (n = 9; 3F, 6M) or a control viral fluorophore (n = 11; 6F, 5M) in the DLS first underwent EtOH-induced stimulant response testing as described above, but only testing locomotor activity following i.p. delivery of saline or 1 g/kg EtOH. This dose was chosen because it is where our first CalEX group showed the greatest stimulant response, and we wanted to limit alcohol exposure prior to home cage binge drinking (‘drinking-in-the-dark’). Then, mice underwent metabolic testing to assess if astrocyte CalEX expression changes EtOH metabolism in mice with the same EtOH history (a single 1 g/kg EtOH injection). For this assessment, all mice received a 2 g/kg injection of EtOH and then 50 µl of retro-orbital blood was drawn 2 hours later. Blood samples were centrifuged, and plasma was withdrawn and stored at -80°C. Blood ethanol concentrations (BECs) were determined using an Analox EtOH Analyzer (Analox Instruments, Lunenburg, MA).

#### CalEX: drinking-in-the-dark and quinine adulterated drinking

Binge-like voluntary EtOH consumption was assessed using a home-cage Drinking-in-the-Dark (DID) paradigm adapted from Rhodes et al. [90]. Mice were housed on a reverse 12:12 h light/dark cycle (lights off at 08:00) to align peak activity with the testing period. To permit accurate measurement of fluid intake, mice were single-housed. Beginning 3 hours into the dark phase (11:00), the home-cage water bottle was replaced with a drinking tube created from a 10mL pipette with attached sipper containing 20% EtOH diluted in tap water (from 190 proof EtOH, Koptec). Mice were given 2 hours of access to the EtOH solution each day (11:00 - 13:00). At the end of each session, EtOH tubes were removed and the standard water bottles were returned. Food was available ad libitum, even during EtOH drinking periods.

EtOH consumption was determined by reading the volume of the drinking tubes immediately before and after each session. Raw consumption was calculated as the difference between initial and final bottle volume. Spillage was controlled by using control sippers placed on empty cages and subtracted from each animal’s measured consumption. Intake was normalized to body weight and expressed as g/kg. DID testing occurred consecutively for 15 days.

To assess aversion resistant alcohol consumption, quinine adulteration tests were conducted during the final phase of the drinking paradigm. On Day 16, mice received 20% EtOH with quinine (250 µM), followed by a washout day (Day 17, 20% EtOH only), an EtOH test day (Day 18, 20% EtOH), a second quinine session (Day 19, 500 µM quinine in 20% EtOH), lastly, on day 20, mice were tested with 250 µM quinine in water only to assess if both viral groups can equally taste the quinine. Quinine hemisulfate salt monohydrate (Sigma-Aldrich Q1250) was dissolved directly into the EtOH or water solution to achieve the desired final concentrations. For assessment of aversion-resistant drinking, day 16 250 µM quinine-adulterated alcohol (QuA) consumption was compared to the final day of non-adulterated DID intake (day 15). For day 19 (500 µM QuA) assessment, intakes were compared to the day 18 EtOH test day.

#### CalEX: loss of righting reflex (LORR)

To assess the role of striatal astrocyte calcium signaling in the sedative effects of alcohol, CalEX and control mice were subsequently assessed for loss of righting reflex (LORR). LORR is a well-established protocol [91] where mice received a 3.5 g/kg intraperitoneal injection of EtOH. For LORR testing, 190 proof EtOH (Koptec) diluted to 45% EtOH in sterile saline was used. Following injection, mice were continuously monitored for the onset of LORR. Latency to LORR was defined as the time from EtOH injection to the point at which the mouse was unable to right itself when placed on its back for 30 seconds. Once LORR was established, mice were placed on v-shaped plexiglass troughs and remained in the trough until recovery of righting reflex. Recovery of the righting reflex was defined as the ability of the mouse to independently right itself three consecutive times within a 30-second period after being placed on its back. LORR duration was defined as the time interval between loss of the righting reflex and recovery of the righting reflex. Total test time was defined as the time from EtOH injection to recovery of the righting reflex.

### Extended Results

#### DLS astrocyte calcium inhibition increased early 2-bottle choice home-cage drinking

We were also interested in assessing the role of DLS astrocyte calcium signaling in modulating other alcohol-related behaviors. For CalEX experiment 2, we assessed the role of striatal astrocytes in modulating home-cage intermittent 2-bottle choice (2BC) alcohol drinking in automated Sable cages (Figure S4A). We found that across the total 23 days of intermittent access to 20% EtOH in a 2BC home-cage drinking paradigm, there were no significant differences in total EtOH intake between CalEX and control groups (Figure S4B). There were no significant differences in other measures, including water intake, EtOH preference, and distance travelled (Figures S4C, D, and E respectively). Of note, there were no main or interaction effects of sex in total EtOH intake (Figure S4B), but there was a main effect of sex for water intake (Figure S4C) and distance (Figure S4E), where females of both viral groups generally consumed more water and moved more than males. There were no interactions of sex with day, virus, or day x virus for these measures (Table 3). The increased female water intake led to a trend for a main effect of sex in EtOH preference, where females overall tended to prefer EtOH more (*p* = 0.0509; Figure S4D). Overall, these results indicated that DLS astrocytes did not regulate voluntary home-cage alcohol drinking or impact other behaviors, such as water intake or total home cage locomotor activity. As a note, the 2BC experiments were performed mice consumed slightly less EtOH than would be expected in a “traditional” 2BC home-cage test in the Sable cages. In the Sable cages, across 23 days (intermittent access), CalEX mice consumed an average of 7.45 ± 0.739 g/kg (mean ± SD) and control mice consumed 6.21 ± 1.23 g/kg (see also Figure S4B). In the subsequent “traditional” 2BC home cage test across 10 days (intermittent access), CalEX mice consumed 14.22 ± 1.19 g/kg and control mice consumed 16.12 ± 1.10 g/kg. No significant main effects of virus or day x virus interactions were present [data not shown].

#### DLS astrocyte calcium inhibition did not impact metabolic rate, home-cage binge drinking, aversion-resistant drinking, or LORR

For the final CalEX experiment, CalEX and control mice underwent a sequence of EtOH-related behavioral testing (Figure S5A). We assayed the stimulation response again in this cohort but tested only a single dose to minimize alcohol exposure for the subsequent DID drinking experiments. We note that this experimental design is not ideal given our previous finding that mice show considerable variability in their maximally stimulating dose (Figure S3B) and that CalEX expression potentiates stimulation mostly at submaximal doses (Figure 2G). Analysis of EtOH-induced stimulant response testing for saline (Figure S5B) and 1 g/kg EtOH (Figure S5C) indicated a main effect of sex for both injections, where females expressing CalEX or a control virus moved more than males, an effect that was not observed in the previous CalEX alcohol stimulation experiment (Figure 2). One possible explanation for the lack of virus effect in this cohort is that, on average, the 1 g/kg dose is not a submaximal dose for a significant proportion of the mice. In this case, any effect present at the individual mouse level may have been lost at the group level. We also assessed alcohol metabolism across the two viral groups. BEC measurements 2 hours after i.p. injection of 2 g/kg alcohol showed no difference between viral groups, indicating that CalEX and control mice metabolize EtOH at a similar rate (Figure S5D).

To further assess the impact of DLS astrocyte calcium inhibition on home-cage drinking, we used another home-cage drinking paradigm focused on binge EtOH intake. Across 15 days of consecutive drinking-in-the-dark (DID) exposure, CalEX and control mice consumed similar amounts of EtOH (Figure S5E). We then tested if DLS astrocyte calcium inhibition impacted aversion-resistant EtOH intake. Mice of either viral group decreased intake when the alcohol was adulterated with 250 µM (Figure S6F) and 500 µM quinine (Figure S5G), indicating that neither viral group displays aversion-resistant alcohol drinking. Analysis of 500 µM quinine-adulterated EtOH intake indicated a significant interaction between solution, sex and virus, *F*(1, 16) = 6.835, *p* = 0.018, η^2^p = 0.299, which was driven primarily by CalEX females consuming high levels unadulterated EtOH while consuming the least amount of quinine-adulterated EtOH of all the groups (CalEX (n = 9; 3F, 6M)). Follow-up on this effect revealed no significant main or interactions of sex with virus or solution (Table 3), therefore, results are shown collapsed on sex. Lastly, we assessed intake of 250 µM quinine-adulterated water which indicated that viral groups can taste the quinine equally (Figure S5H).

Lastly, we assessed alcohol sedation to an i.p. injection of 3.5g/kg EtOH by performing the loss of righting reflex (LORR) assay. We identified an interesting main effect of sex in latency to LORR (Figure S5I), where male mice took longer to lose righting reflex than female mice. Of note, two CalEX male mice did not loss their righting reflex for > 1 hour following the 3.5 g/kg EtOH injection and were omitted from this analysis. This phenomenon was not observed in any other group. We did not observe differences between viral groups in any LORR dependent variables (Figures S5J-L). These results indicate that our previous finding that DLS astrocyte calcium inhibition increased EtOH-induced stimulation (Figure 2) is specific to stimulation and does not appear to be related to EtOH-induced sedation.

## Notes

### Competing Interest Statement

The authors have declared no competing interest.

## References

1. Stokłosa, I., et al., Medications for the Treatment of Alcohol Dependence-Current State of Knowledge and Future Perspectives from a Public Health Perspective. Int J Environ Res Public Health, 2023. 20(3).

2. Adermark, L. and M.S. Bowers, Disentangling the Role of Astrocytes in Alcohol Use Disorder. Alcohol Clin Exp Res, 2016. 40(9): p. 1802–16.

3. Parpura, V., et al., Glutamate-mediated astrocyte-neuron signalling. Nature, 1994. 369(6483): p. 744-7.

4. Wang, F., et al., Astrocytes modulate neural network activity by Ca²+-dependent uptake of extracellular K+. Sci Signal, 2012. 5(218): p. ra26.

5. Nowacka, A., M. Śniegocki, and E.A. Ziółkowska, Calcium Dynamics in Astrocyte-Neuron Communication from Intracellular to Extracellular Signaling. Cells, 2025. 14(21).

6. Araque, A., et al., Tripartite synapses: glia, the unacknowledged partner. Trends Neurosci, 1999. 22(5): p. 208–15.

7. Covelo, A. and A. Araque, Lateral regulation of synaptic transmission by astrocytes. Neuroscience, 2016. 323: p. 62–6.

8. Lines, J., et al., Astrocytes modulate sensory-evoked neuronal network activity. Nature Communications, 2020. 11(1): p. 3689.

9. Purushotham, S.S. and Y. Buskila, Astrocytic modulation of neuronal signalling. Front Netw Physiol, 2023. 3: p. 1205544.

10. Oliveira, J.F. and A. Araque, Astrocyte regulation of neural circuit activity and network states. Glia, 2022. 70(8): p. 1455–1466.

11. Erickson, E.K., et al., Cortical astrocytes regulate ethanol consumption and intoxication in mice. Neuropsychopharmacology, 2021. 46(3): p. 500–508.

12. Nwachukwu, K.N., et al., Chemogenetic manipulation of astrocytic signaling in the basolateral amygdala reduces binge-like alcohol consumption in male mice. J Neurosci Res, 2021. 99(8): p. 1957–1972.

13. Giacometti, L.L., et al., Astrocyte modulation of extinction impairments in ethanol-dependent female mice. Neuropharmacology, 2020. 179: p. 108272.

14. Holdstock, L. and H. de Wit, Individual differences in the biphasic effects of ethanol. Alcohol Clin Exp Res, 1998. 22(9): p. 1903–11.

15. King, A.C., et al., Biphasic alcohol response differs in heavy versus light drinkers. Alcohol Clin Exp Res, 2002. 26(6): p. 827–35.

16. Erblich, J. and M. Earleywine, Behavioral undercontrol and subjective stimulant and sedative effects of alcohol intoxication: independent predictors of drinking habits? Alcohol Clin Exp Res, 2003. 27(1): p. 44–50.

17. Gmel, G., et al., Association of anticipated stimulant and sedative effects of alcohol with future heavy drinking in a large Swiss cohort study of young men. Journal of Studies on Alcohol and Drugs. 0(ja): p. jsad.25-00189.

18. King, A.C., et al., Rewarding, stimulant, and sedative alcohol responses and relationship to future binge drinking. Arch Gen Psychiatry, 2011. 68(4): p. 389–99.

19. King, A.C., et al., Alcohol challenge responses predict future alcohol use disorder symptoms: a 6-year prospective study. Biol Psychiatry, 2014. 75(10): p. 798–806.

20. King, A., et al., Subjective Responses to Alcohol in the Development and Maintenance of Alcohol Use Disorder. Am J Psychiatry, 2021. 178(6): p. 560–571.

21. King, A.C., et al., The role of alcohol response phenotypes in the risk for alcohol use disorder. BJPsych Open, 2019. 5(3): p. e38.

22. Wise, R.A. and M.A. Bozarth, A psychomotor stimulant theory of addiction. Psychol Rev, 1987. 94(4): p. 469–92.

23. Newlin, D.B. and J.B. Thomson, Alcohol challenge with sons of alcoholics: a critical review and analysis. Psychol Bull, 1990. 108(3): p. 383–402.

24. Belnap, M.A., et al., Subjective response to alcohol predicts motivation to self-administer alcohol in a progressive ratio task. Experimental and Clinical Psychopharmacology, 2025. 33(5): p. 503–512.

25. Bocarsly, M.E., et al., A Mechanism Linking Two Known Vulnerability Factors for Alcohol Abuse: Heightened Alcohol Stimulation and Low Striatal Dopamine D2 Receptors. Cell Reports, 2019. 29(5): p. 1147–1163.e5.

26. Lessov, C.N., et al., Voluntary ethanol drinking in C57BL/6J and DBA/2J mice before and after sensitization to the locomotor stimulant effects of ethanol. Psychopharmacology (Berl), 2001. 155(1): p. 91–9.

27. Boehm, S.L., 2nd, et al., Ventral tegmental area region governs GABA(B) receptor modulation of ethanol-stimulated activity in mice. Neuroscience, 2002. 115(1): p. 185–200.

28. Linsenbardt, D.N. and S.L. Boehm, 2nd, Determining the heritability of ethanol-induced locomotor sensitization in mice using short-term behavioral selection. Psychopharmacology (Berl), 2013. 230(2): p. 267–78.

29. Spuhler, K. and R.A. Deitrich, Correlative analysis of ethanol-related phenotypes in rat inbred strains. Alcohol Clin Exp Res, 1984. 8(5): p. 480–4.

30. Weafer, J., et al., Striatal activity correlates with stimulant-like effects of alcohol in healthy volunteers. Neuropsychopharmacology, 2018. 43(13): p. 2532–2538.

31. Nakano, K., et al., Neural circuits and functional organization of the striatum. Journal of Neurology, 2000. 247(5): p. V1–V15.

32. Pakhotin, P. and E. Bracci, Cholinergic Interneurons Control the Excitatory Input to the Striatum. The Journal of Neuroscience, 2007. 27(2): p. 391–400.

33. Cox, J. and I.B. Witten, Striatal circuits for reward learning and decision-making. Nature Reviews Neuroscience, 2019. 20(8): p. 482–494.

34. Ye, L., et al., Ethanol abolishes vigilance-dependent astroglia network activation in mice by inhibiting norepinephrine release. Nat Commun, 2020. 11(1): p. 6157.

35. Catlin, M.C., T.J. Kavanagh, and L.G. Costa, Muscarinic receptor-induced calcium responses in astroglia. Cytometry, 2000. 41(2): p. 123–32.

36. Shen, J.-x. and J.L. Yakel, Nicotinic acetylcholine receptor-mediated calcium signaling in the nervous system. Acta Pharmacologica Sinica, 2009. 30(6): p. 673–680.

37. Murphy, S., B. Pearce, and C. Morrow, Astrocytes have both M1 and M2 muscarinic receptor subtypes. Brain Research, 1986. 364(1): p. 177–180.

38. Navarrete, M., et al., Astrocytes mediate in vivo cholinergic-induced synaptic plasticity. PLoS Biol, 2012. 10(2): p. e1001259.

39. Kalelkar, A., et al., A paradigm for ethanol consumption in head-fixed mice during prefrontal cortical two-photon calcium imaging. Neuropharmacology, 2024. 245: p. 109800.

40. Evans, W.R., et al., Chemogenetic Control of Striatal Astrocytes Improves Parkinsonian Motor Deficits in Mice. Glia, 2025. 73(6): p. 1188–1202.

41. Ambrosi, P. and T.N. Lerner, Striatonigrostriatal circuit architecture for disinhibition of dopamine signaling. Cell Rep, 2022. 40(7): p. 111228.

42. Mathis, A., et al., DeepLabCut: markerless pose estimation of user-defined body parts with deep learning. Nature Neuroscience, 2018. 21(9): p. 1281–1289.

43. Yu, X., et al., Reducing Astrocyte Calcium Signaling In Vivo Alters Striatal Microcircuits and Causes Repetitive Behavior. Neuron, 2018. 99(6): p. 1170–1187.e9.

44. Kim, E., et al., Prefrontal cortex astrocytes modulate distinct neuronal populations to control anxiety-like behavior. Nature Communications, 2025. 16(1): p. 7819.

45. Bukalo, O., et al., Astrocytes enable amygdala neural representations supporting memory. Nature, 2026.

46. Yu, X., S.L. Moye, and B.S. Khakh, Local and CNS-Wide Astrocyte Intracellular Calcium Signaling Attenuation In Vivo with CalEx(flox) Mice. J Neurosci, 2021. 41(21): p. 4556–4574.

47. Peyton, L., et al., In vivo calcium extrusion from accumbal astrocytes reduces anxiety-like behaviors but increases compulsive-like responses and compulsive ethanol drinking in mice. Neuropharmacology, 2025. 268: p. 110320.

48. Luchsinger, J.R., et al., Delineation of an insula-BNST circuit engaged by struggling behavior that regulates avoidance in mice. Nat Commun, 2021. 12(1): p. 3561.

49. Flavin, S.A. and D.G. Winder, Noradrenergic control of the bed nucleus of the stria terminalis in stress and reward. Neuropharmacology, 2013. 70: p. 324–30.

50. Feng, J., et al., A Genetically Encoded Fluorescent Sensor for Rapid and Specific In Vivo Detection of Norepinephrine. Neuron, 2019. 102(4): p. 745–761.e8.

51. Huang, Z., et al., Dynamic responses of striatal cholinergic interneurons control behavioral flexibility. Science Advances, 2024. 10(51): p. eadn2446.

52. Middauch, L.D., W.O. Boggan, and C.L. Randall, Stimulatory effects of ethanol in C57BL/6 mice. Pharmacology Biochemistry and Behavior, 1987. 27(3): p. 421–424.

53. Crabbe, J.C., et al., Mice genetically selected for differences in open-field activity after ethanol. Pharmacol Biochem Behav, 1987. 27(3): p. 577–81.

54. Allen, N.J. and C. Eroglu, Cell Biology of Astrocyte-Synapse Interactions. Neuron, 2017. 96(3): p. 697–708.

55. Lines, J., et al., The Duality of Astrocyte Neuromodulation: Astrocytes Sense Neuromodulators and Are Neuromodulators. Journal of Neurochemistry, 2025. 169(4): p. e70054.

56. Porter, J.T. and K.D. McCarthy, Astrocytic neurotransmitter receptors in situ and in vivo. Prog Neurobiol, 1997. 51(4): p. 439–55.

57. Agulhon, C., et al., What is the role of astrocyte calcium in neurophysiology? Neuron, 2008. 59(6): p. 932–46.

58. Nagai, J., et al., Hyperactivity with Disrupted Attention by Activation of an Astrocyte Synaptogenic Cue. Cell, 2019. 177(5): p. 1280–1292.e20.

59. López, R.C., et al., Innervation density governs crosstalk of GPCR-based norepinephrine and dopamine sensors. bioRxiv, 2024.

60. Feng, J., et al., Monitoring norepinephrine release in vivo using next-generation GRABNE sensors. Neuron, 2024. 112(12): p. 1930–1942.e6.

61. Butcher, S.G. and L.L. Butcher, Origin and modulation of acetylcholine activity in the neostriatum. Brain Research, 1974. 71(1): p. 167–171.

62. Stedehouder, J., et al., Rapid modulation of striatal cholinergic interneurons and dopamine release by satellite astrocytes. bioRxiv, 2024: p. 2024.05.15.594341.

63. Slade, L.E., et al., Neurotoxic effects of chronic ethanol on cholinergic interneurons in the dorsomedial striatum. Neuropharmacology, 2026. 282: p. 110706.

64. Fedotova, A., et al., Dissociation Between Neuronal and Astrocytic Calcium Activity in Response to Locomotion in Mice. Function (Oxf), 2023. 4(4): p. zqad019.

65. Dombeck, D.A., et al., Imaging large-scale neural activity with cellular resolution in awake, mobile mice. Neuron, 2007. 56(1): p. 43–57.

66. Reitman, M.E., et al., Norepinephrine links astrocytic activity to regulation of cortical state. Nature Neuroscience, 2023. 26(4): p. 579–593.

67. Åbjørsbråten, K.S., et al., Impaired astrocytic Ca2+ signaling in awake-behaving Alzheimer’s disease transgenic mice. eLife, 2022. 11: p. e75055.

68. Scibelli, A.C. and T.J. Phillips, Combined scopolamine and ethanol treatment results in a locomotor stimulant response suggestive of synergism that is not blocked by dopamine receptor antagonists. Alcohol Clin Exp Res, 2009. 33(3): p. 435–47.

69. Chen, G., et al., Striatal involvement in human alcoholism and alcohol consumption, and withdrawal in animal models. Alcohol Clin Exp Res, 2011. 35(10): p. 1739–48.

70. Everitt, B.J. and T.W. Robbins, Neural systems of reinforcement for drug addiction: from actions to habits to compulsion. Nat Neurosci, 2005. 8(11): p. 1481–9.

71. Wilcox, M.V., et al., Repeated binge-like ethanol drinking alters ethanol drinking patterns and depresses striatal GABAergic transmission. Neuropsychopharmacology, 2014. 39(3): p. 579–94.

72. DePoy, L., et al., Chronic alcohol produces neuroadaptations to prime dorsal striatal learning. Proceedings of the National Academy of Sciences, 2013. 110(36): p. 14783–14788.

73. Koob, G.F. and N.D. Volkow, Neurocircuitry of addiction. Neuropsychopharmacology, 2010. 35(1): p. 217–38.

74. Hwang, S.N., et al., Astrocytic Regulation of Neural Circuits Underlying Behaviors. Cells, 2021. 10(2).

75. Huang, M., et al., Astrocytes in the medial entorhinal cortex are required for spatial exploration. Cell Rep, 2025. 44(9): p. 116173.

76. Vena, A.A. and R.A. Gonzales, Temporal profiles dissociate regional extracellular ethanol versus dopamine concentrations. ACS Chem Neurosci, 2015. 6(1): p. 37–47.

77. Nielsen, B.E. and C.P. Ford, Reduced striatal M4-cholinergic signaling following dopamine loss contributes to parkinsonian and l-DOPA-induced dyskinetic behaviors. Sci Adv, 2024. 10(47): p. eadp6301.

78. Hansson, E. and L. Rönnbäck, *Astrocytic receptors and second messenger systems*, in Advances in Molecular and Cell Biology. 2003, Elsevier. p. 475–501.

79. Wang, X., et al., Activation of α7-containing nicotinic receptors on astrocytes triggers AMPA receptor recruitment to glutamatergic synapses. J Neurochem, 2013. 127(5): p. 632–43.

80. Van Der Zee, E.A., et al., Muscarinic acetylcholine receptor-expression in astrocytes in the cortex of young and aged rats. Glia, 1993. 8(1): p. 42–50.

81. Li, W.-P., et al., Astrocytes Mediate Cholinergic Regulation of Adult Hippocampal Neurogenesis and Memory Through M1 Muscarinic Receptor. Biological Psychiatry, 2022. 92(12): p. 984–998.

82. Araque, A., et al., Synaptically released acetylcholine evokes Ca2+ elevations in astrocytes in hippocampal slices. J Neurosci, 2002. 22(7): p. 2443–50.

83. Takata, N., et al., Astrocyte calcium signaling transforms cholinergic modulation to cortical plasticity in vivo. J Neurosci, 2011. 31(49): p. 18155–65.

84. Aosaki, T., A.M. Graybiel, and M. Kimura, Effect of the nigrostriatal dopamine system on acquired neural responses in the striatum of behaving monkeys. Science, 1994. 265(5170): p. 412-5.

85. Di Chiara, G. and A. Imperato, Ethanol preferentially stimulates dopamine release in the nucleus accumbens of freely moving rats. Eur J Pharmacol, 1985. 115(1): p. 131–2.

86. Krok, A.C., et al., Intrinsic dopamine and acetylcholine dynamics in the striatum of mice. Nature, 2023. 621(7979): p. 543–549.

87. Chantranupong, L., et al., Dopamine and glutamate regulate striatal acetylcholine in decision-making. Nature, 2023. 621(7979): p. 577–585.

88. Guttenplan, K.A., et al., GPCR signaling gates astrocyte responsiveness to neurotransmitters and control of neuronal activity. Science, 2025. 388(6748): p. 763–768.

89. Wise, R.A., Voluntary ethanol intake in rats following exposure to ethanol on various schedules. Psychopharmacologia, 1973. 29(3): p. 203–10.

90. Rhodes, J.S., et al., Evaluation of a simple model of ethanol drinking to intoxication in C57BL/6J mice. Physiol Behav, 2005. 84(1): p. 53–63.

91. Erwin, V.G., V.M. Gehle, and R.A. Deitrich, Selectively bred lines of mice show response and drug specificity for genetic regulation of acute functional tolerance to ethanol and pentobarbital. J Pharmacol Exp Ther, 2000. 293(1): p. 188–95.

